# “CapZyme-Seq” comprehensively defines promoter-sequence determinants for RNA 5’ capping with NAD^+^

**DOI:** 10.1101/239426

**Authors:** Irina O. Vvedenskaya, Jeremy G. Bird, Yuanchao Zhang, Yu Zhang, Xinfu Jiao, Ivan Barvík, Libor Krásný, Megerditch Kiledjian, Deanne M. Taylor, Richard H. Ebright, Bryce E. Nickels

## Abstract

Nucleoside-containing metabolites such as NAD^+^ can be incorporated as “5′ caps” on RNA by serving as non-canonical initiating nucleotides (NCINs) for transcription initiation by RNA polymerase (RNAP). Here, we report “CapZyme-Seq,” a high-throughput-sequencing method that employs NCIN-decapping enzymes NudC and Rai1 to detect and quantify NCIN-capped RNA. By combining CapZyme-Seq with multiplexed transcriptomics, we determine efficiencies of NAD^+^ capping by *Escherichia coli* RNAP for ~16,000 promoter sequences. The results define preferred transcription start-site (TSS) positions for NAD^+^ capping and define a consensus promoter sequence for NAD^+^ capping: HRRASWW (TSS underlined). By applying CapZyme-Seq to *E. coli* total cellular RNA, we establish that sequence determinants for NCIN capping *in vivo* match the NAD^+^-capping consensus defined *in vitro*, and we identify and quantify NCIN-capped small RNAs. Our findings define the promoter-sequence determinants for NCIN capping with NAD^+^ and provide a general method for analysis of NCIN capping *in vitro* and *in vivo*.

## INTRODUCTION

RNA 5′-end capping provides a layer of “epitranscriptomic” regulation (Jaschke et al., 2016). RNA 5′-end capping influences RNA fate, including stability, processing, localization and translatability; and RNA capping enables cells to distinguish between host and pathogen RNA (Topisirovic et al., 2011; Jaschke et al., 2016; Ramanathan et al., 2016). Cellular processes that add, remove, or modify RNA caps modulate RNA fate and the distinction between host and pathogen RNA (Li and Kiledjian, 2010; Arribas-Layton et al., 2013; Jaschke et al., 2016; Ramanathan et al., 2016; Grudzien-Nogalska and Kiledjian, 2017).

One form of RNA 5′-capping, which has been observed solely in eukaryotes and certain eukaryotic viruses (Wei et al., 1975; Shatkin, 1976; Furuichi and Shatkin, 2000), entails addition of a 7-methylguanylate (m^7^G) to the RNA 5′ end. The m^7^G cap is added to the nascent RNA after transcription initiation, when the nascent RNA reaches a length of ~20 nt, and is added by a capping complex that interacts with the nascent RNA transcript as it emerges from RNA polymerase (RNAP) (Shuman, 2001; Ghosh and Lima, 2010; Martinez-Rucobo et al., 2015; Shuman, 2015). The m^7^G cap protects RNA from exonuclease digestion, allows cells to distinguish ‘self’ and ‘foreign’ RNA, and enables recognition and binding of proteins that facilitate splicing, polyadenylation, nuclear export, and translation efficiency (Topisirovic et al., 2011; Devarkar et al., 2016; Ramanathan et al., 2016).

Recently, a second form of RNA 5′-capping has been identified, first in bacteria (Chen et al., 2009; Kowtoniuk et al., 2009; Cahova et al., 2015; Bird et al., 2016), and then in eukaryotes (Jiao et al., 2017; Walters et al., 2017). In this second form of RNA capping, a nucleoside-containing metabolite such as nicotinamide adenine dinucleotide (NAD^+^) is added at the RNA 5′ end. The nucleoside-containing metabolite cap is introduced during the first nucleotide addition step in transcription initiation, and is added by RNAP itself (Bird et al., 2016; Hofer and Jaschke, 2016; Barvik et al., 2017). The nucleoside-containing metabolite serves as a “non-canonical initiating nucleotide” (NCIN) for transcription initiation by RNAP, providing an *“ab initio*” mechanism of RNA capping (Bird et al., 2016). As with m^7^G caps, NCIN caps modulate RNA fate by influencing RNA stability and translation efficiency (Cahova et al., 2015; Bird et al., 2016; Jiao et al., 2017). NCIN capping has been observed *in vitro* with both bacterial RNAP (Malygin and Shemyakin, 1979; Bird et al., 2016; Julius and Yuzenkova, 2017) and eukaryotic RNAP II (Bird et al., 2016). NCIN-capped RNAs have been observed *in vivo* for bacteria (Chen et al., 2009; Kowtoniuk et al., 2009; Cahova et al., 2015; Bird et al., 2016; Nubel et al., 2017), yeast (Walters et al., 2017), and human cells in culture (Jiao et al., 2017), and the *ab initio* mechanism of RNA-capping with NCINs has been demonstrated *in vivo* in bacteria (Bird et al., 2016).

In addition to NAD^+^, other nucleoside-containing metabolites, including reduced NAD^+^ (NADH), 5′-desphospho coenzyme A (dpCoA), flavin adenine dinucleotide (FAD), uridine diphosphate glucose (UDP-glucose), and uridine diphosphate N-acetylglucosamine (UDP-GlcNAc), can serve as substrates for NCIN-mediated capping *in vitro* (Malygin and Shemyakin, 1979; Bird et al., 2016; Julius and Yuzenkova, 2017), suggesting that these nucleoside-containing metabolites also potentially could function in NCIN capping *in vivo*.

Adenosine-containing NCINs (NAD^+^, NADH, dpCoA, and FAD; Figure 1A) compete with adenosine triphosphate (ATP) for use by RNAP as initiating nucleotides. Uridine-containing NCINs (UDP-glucose and UDP-glcNAc) compete with uridine triphosphate (UTP) for use by RNAP as initiating nucleotides. We have presented evidence that promoter sequence at and immediately upstream of the transcription start site determines the outcome of the competition between initiation with NCINs and initiation with NTPs (Bird et al., 2016). Julius and Yuzenkova have challenged this evidence (Julius and Yuzenkova, 2017). Here, we report “CapZyme-Seq,” a next-generation-sequencing-based method for detection and quantitation of NCIN-capped RNAs. We apply this method to define, comprehensively, the promoter-sequence determinants and promoter-consensus sequence for NCIN capping with NAD^+^ *in vitro,* to define the promoter-sequence determinants and promoter-consensus sequence for NCIN capping *in vivo,* and to identify and quantify NCIN-capped small RNAs *in vivo*.

**Figure 1.**
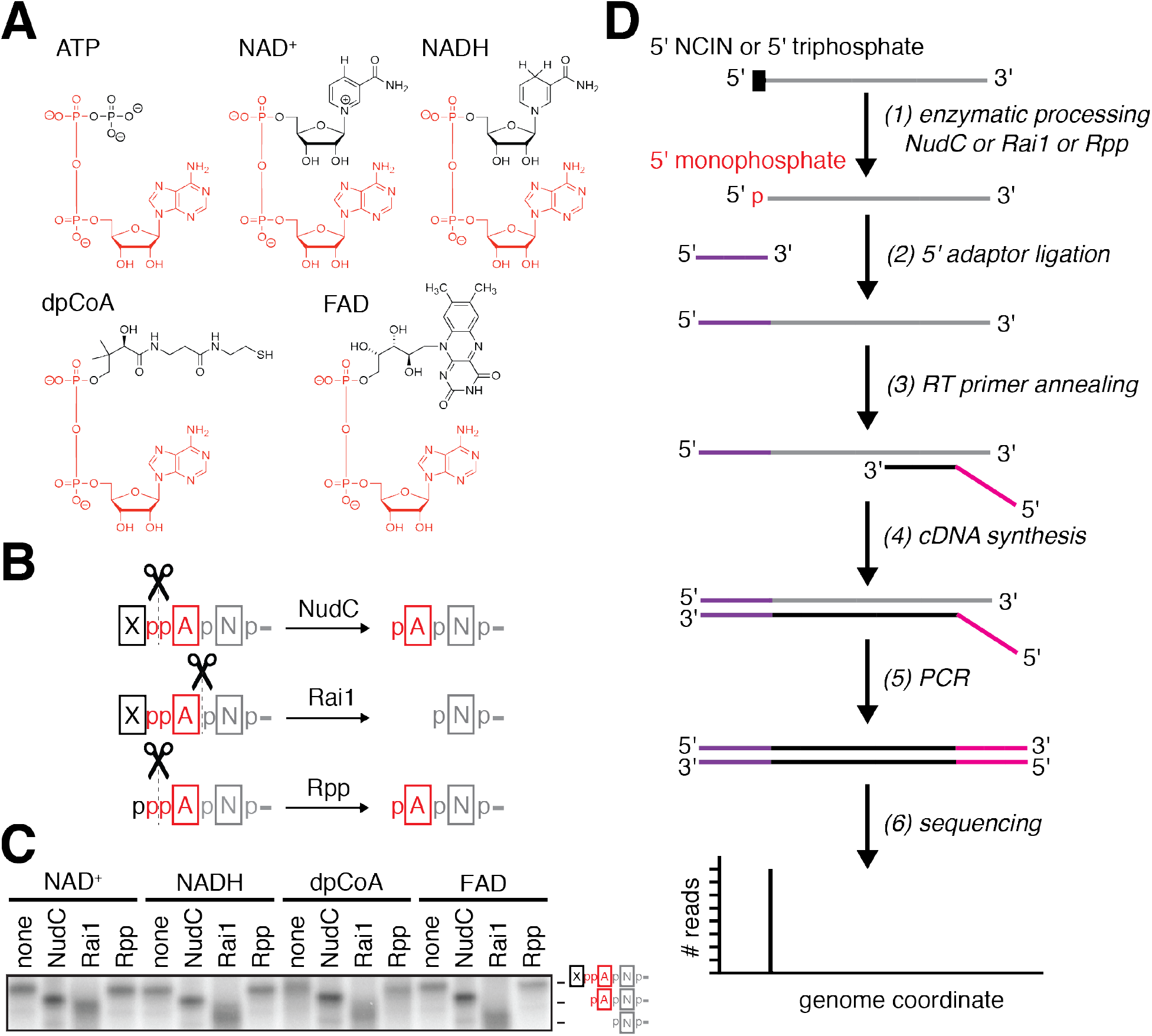
CapZyme-Seq, a high-throughput-sequencing method to detect NCIN-capped RNA. **A**. Structures of ATP and adenosine-containing NCINs NAD^+^, NADH, dpCoA, and FAD. Red, identical atoms. **B**. Processing of RNA 5′-ends by NudC, Rai1, and Rpp. Red, common moiety of each 5′-end; black, distinct moiety of each 5′-end; grey, remainder of RNA. **C**. Products of processing of NCIN-capped RNA 5′-ends by NudC, Rai1, and Rpp. **D**. CapZyme-Seq procedure. Grey, RNA; purple, 5′ adaptor; red, 3′ adaptor; black cDNA.

## RESULTS AND DISCUSSION

### CapZyme-Seq, a high-throughput-sequencing method to detect NCIN-capped RNA

Methods for high-throughput sequencing of RNA 5′-ends often rely on the ligation of singlestranded oligonucleotide adaptors to RNA 5′-ends. Adaptor ligation requires that the RNA have a 5′-monophosphate. Accordingly, analysis of RNAs that do not have a 5′-monophosphate requires enzymatic processing of the RNAs to yield RNAs having a 5′-monophosphate. Here, we exploit the requirement for enzymatic processing to yield a 5′-monophosphate, together with the use of processing enzymes specific for NCIN-capped RNA and a processing enzyme specific for uncapped, 5’-triphosphate RNA, to enable differential detection and quantitation of NCIN-capped RNA and uncapped, 5’-triphosphate RNA. We term this method “CapZyme-Seq” (Figure 1).

For selective processing of NCIN-capped RNA to 5′-monophosphate RNA, we used the bacterial RNA-decapping enzyme NudC or the fungal RNA-decapping enzyme Rai1. NudC processes NCIN-capped RNA to 5′-monophosphate RNA by cleaving the diphosphate group of the NCIN cap (Cahova et al., 2015; Hofer et al., 2016), yielding products comprising 5′-pNp-(where N is the 3′ nucleoside moiety of the NCIN, for example the adenosine moiety of NAD^+^) followed by the remainder of the RNA (Figure 1B). Rai1 processes NCIN-capped RNAs to 5′-monophosphate RNA by cleaving the phosphodiester bond connecting the NCIN cap to the remainder of the RNA (Jiao et al., 2017), yielding products comprising 5′-p-followed by the remainder of the RNA (Figure 1B). In initial work, we have found that NudC and Rai1 process RNA capped with at least four of the nucleoside-containing metabolites that can serve as NCINs: NAD^+^, NADH, dpCoA, and FAD (Figure 1C). Accordingly, we propose that CapZyme-Seq using NudC or Rai1 (Figure 1D) can be used to detect NAD^+^-, NADH-, dpCoA-, and FAD-capped RNA.

For selective processing of uncapped, 5′-triphosphate RNA to 5′-monophosphate RNA we used the RNA processing enzyme RNA polyphosphatase, Rpp. Rpp processes 5′-triphosphate RNA to 5′-monophosphate RNA by cleaving the phosphodiester bond between the triphosphate β and α phosphates, yielding products comprising 5′-p-followed by the remainder of the RNA (Figure 1B). In initial work, we have found that Rpp does not detectably process RNAs having any of the above NCIN caps: NAD^+^, NADH, dpCoA, and FAD (Figure 1C).

### CapZyme-Seq analysis of NCIN capping with NAD^+^ in vitro

To define promoter-sequence determinants for NCIN capping, we combined CapZyme-Seq with a multiplexed-transcriptomics method termed “<u>ma</u>ssively <u>s</u>ystematic <u>t</u>ranscript <u>e</u>nd <u>r</u>eadout” (MASTER; Figure 2A). MASTER enables measurement of RNA 5′-end sequences and RNA yields for RNAs generated during transcription of a template library of up to at least 4^10^ barcoded template sequences (Vvedenskaya et al., 2015; Winkelman et al., 2016). Accordingly, combining CapZyme-Seq with MASTER enables measurement of RNA 5′-end sequences and RNA yields for both NCIN-capped RNA and uncapped, 5′-triphosphate RNA for each of up to at least 4^10^ promoter sequences (Figure 2A).

**Figure 2.**
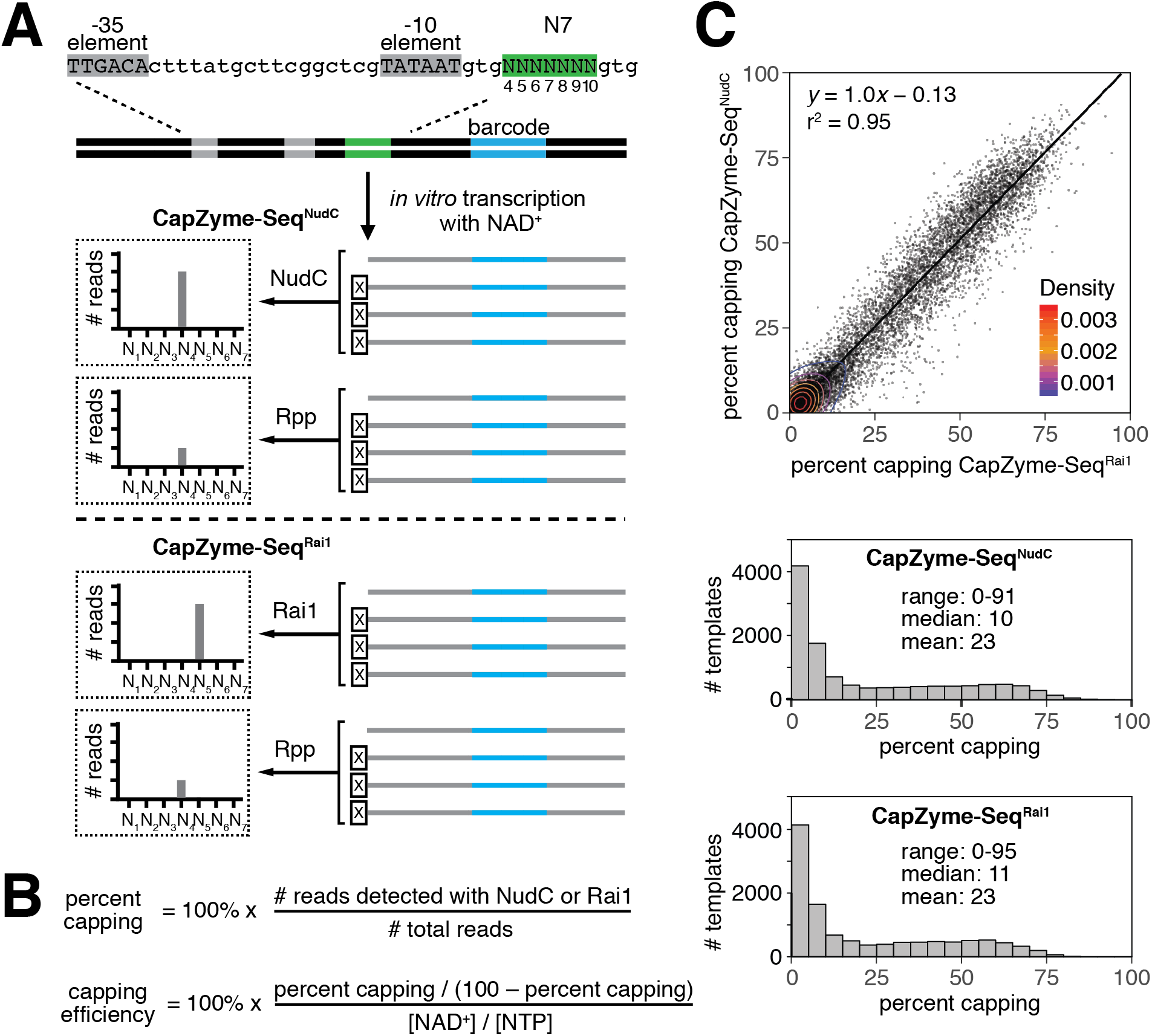
CapZyme-Seq analysis of NCIN capping with NAD^+^ *in vitro*. **A**. Use of CapZyme-Seq in combination with massively systematic transcript end readout (MASTER). Top, lacCONS-N7 promoter library (4^7^, ~16,000 promoter sequences). Grey, promoter −35 and −10 elements; green, randomized sequences 4−10 bp downstream of promoter −10 element; blue, transcribed-region barcode. The linear DNA template contains 94 bp of transcribed-region sequence downstream of the green randomized sequence. Thus, RNA products generated from the lacCONS-N7 promoter library are ~100-nt in length. Middle, CapZyme-Seq using NudC for processing of NCIN-capped RNA and Rpp for processing of uncapped 5′-triphosphate RNA (CapZyme-Seq^NudC^). Bottom, CapZyme-Seq using Rai1 for processing of NCIN-capped RNA and Rpp for processing of uncapped 5′-triphosphate RNA (CapZyme-Seq^Rai1^). **B**. Equations used to calculate percent capping and capping efficiencies. **C**. Results of CapZyme-Seq^NudC^ and CapZyme-Seq^Rai1^. Top, mean percent capping from CapZyme-Seq^NudC^ (n=3) vs. mean percent capping from CapZyme-Seq^Rai1^ (n=3) for ~16,000 promoter sequences (Density from Gaussian kernel density estimation method). Bottom, percent capping histograms.

In this work, we used the MASTER promoter library “*lacCONS-N7*” (Vvedenskaya et al., 2015), which contains 4^7^ (~16,000) derivatives of a consensus *E. coli* σ^70^-dependent promoter comprising all possible sequence variants at the positions 4 to 10 base pairs (bp) downstream of the promoter −10 element (positions 4, 5, 6, 7, 8, 9 and 10; Figure 2A, top). We performed *in vitro* transcription experiments using the *lacCONS*-N7 promoter library and *E. coli* RNAP σ^70^ holoenzyme, in parallel, in the presence or absence of NAD^+^. RNA products from each reaction were analyzed with CapZyme-Seq using NudC (CapZyme-Seq^NudC^; Figure 2A, middle) or Rai1 (CapZyme-Seq^Rai1^; Figure 2A, bottom). We determined “percent capping” (capped RNA yields relative to total RNA yields) and “capping efficiency” [NAD^+^-mediated initiation relative to NTP-mediated initiation; (K_cat_/K_m_,NAD^+^)/(K_cat_/K_m_,NTP)] from the resulting RNA yields using the equations in Figure 2B. Comparison of results obtained using CapZyme-Seq^NudC^ with results obtained using CapZyme-Seq^Rai1^ indicates the results are well correlated (r^2^ ~ 0.95; slope ~ 1.0; Figure 2C, top). The mean percent capping observed is ~23%, the median percent capping is ~10%, and the range of percent capping is 0-95% for the 4^7^ promoter sequences (Figure 2C, bottom). The majority of percent capping values fall within the range of 0-15%. The distribution of percent capping values is highly skewed with a high peak of 0-5% and a long tail extending to greater than 90% (Figure 2C, bottom). The skewed, long-tailed distribution of percent capping confirms that different promoter sequences differ markedly in efficiency of NCIN capping with NAD^+^.

### Determinants for transcription start site selection in NCIN capping with NAD^+^ in vitro

In bacterial transcription initiation, RNAP selects a transcription start site (TSS) at a variable distance downstream of the promoter -10 element. In prior work, we used MASTER to analyze TSS selection in NTP-mediated initiation with the *lacCONS-N7* promoter library (Vvedenskaya et al., 2015). Results indicated that TSS selection occurs over a range of five positions located 6-10 bp downstream of the -10 element (positions 6, 7, 8, 9 and 10), that the preferred, modal position for TSS selection is position 7, and that the order of preference for TSS selection is 7 > 8 > 9 > 6 > 10. Results further indicated that there is a strong sequence preference for G or A (R) at each TSS position.

To define, comprehensively, the determinants for TSS selection in NAD^+^-mediated initiation, we used the combination of CapZyme-Seq and MASTER to determine 5′-end sequence and yields of NAD^+^-capped RNA in NAD^+^-mediated initiation with the *lacCONS-N7* promoter library (Figure 3A-B). To compare these determinants with determinants for TSS selection in NTP-mediated initiation under identical reaction conditions, we used the combination of CapZyme-Seq and MASTER to determine 5′-end sequence and yields of uncapped, 5′-triphosphate RNA in NTP-mediated initiation with the *lacCONS-N7* promoter library in the presence or absence of NAD^+^ (Figures 3C and S1). As with the results above for percent capping (Figure 2C), the results here for TSS selection obtained using CapZyme-Seq^NudC^ and CapZyme-Seq^Rai1^ are well correlated (r^2^ ~ 0.95; slope ~ 1.0; Figure 3B,C and Figure S1B, top). The positional preferences for TSS selection in NAD^+^-mediated initiation (range = positions 6-10; mode = 7 bp downstream of -10 element; mean 7.5 bp downstream of -10 element; order of preference = 7 > 8 > 9 > 6 > 10; Figure 3B, middle) are indistinguishable to the positional preferences for TSS selection in NTP-mediated initiation (range = positions 6-10; mode = 7 bp downstream of -10 element; mean 7.6 bp downstream of -10 element; order of preference = 7 > 8 > 9 > 6 > 10 in reactions performed both in the presence or absence of NAD^+^; Figure 3C, middle, and Figure S1B, middle). However, the sequence preferences for TSS selection in NAD^+^-mediated initiation differ from the sequence preferences for TSS selection in NTP-mediated initiation, exhibiting an essentially absolute preference for TSS positions where the base pair is A:T (Figure 3B, bottom), instead of preference for TSS positions where the base pair is either A:T or G:C (Figure 3C, bottom), consistent with expectation based on the base pairing preferences of the adenosine moiety of NAD^+^ (Figure 1A). Furthermore, in the presence of NAD^+^, there is a decrease in ATP-mediated initiation, but not GTP-mediated initiation (Figure S1B, bottom), consistent with competition between initiation with NAD^+^ and initiation with ATP.

**Figure 3.**
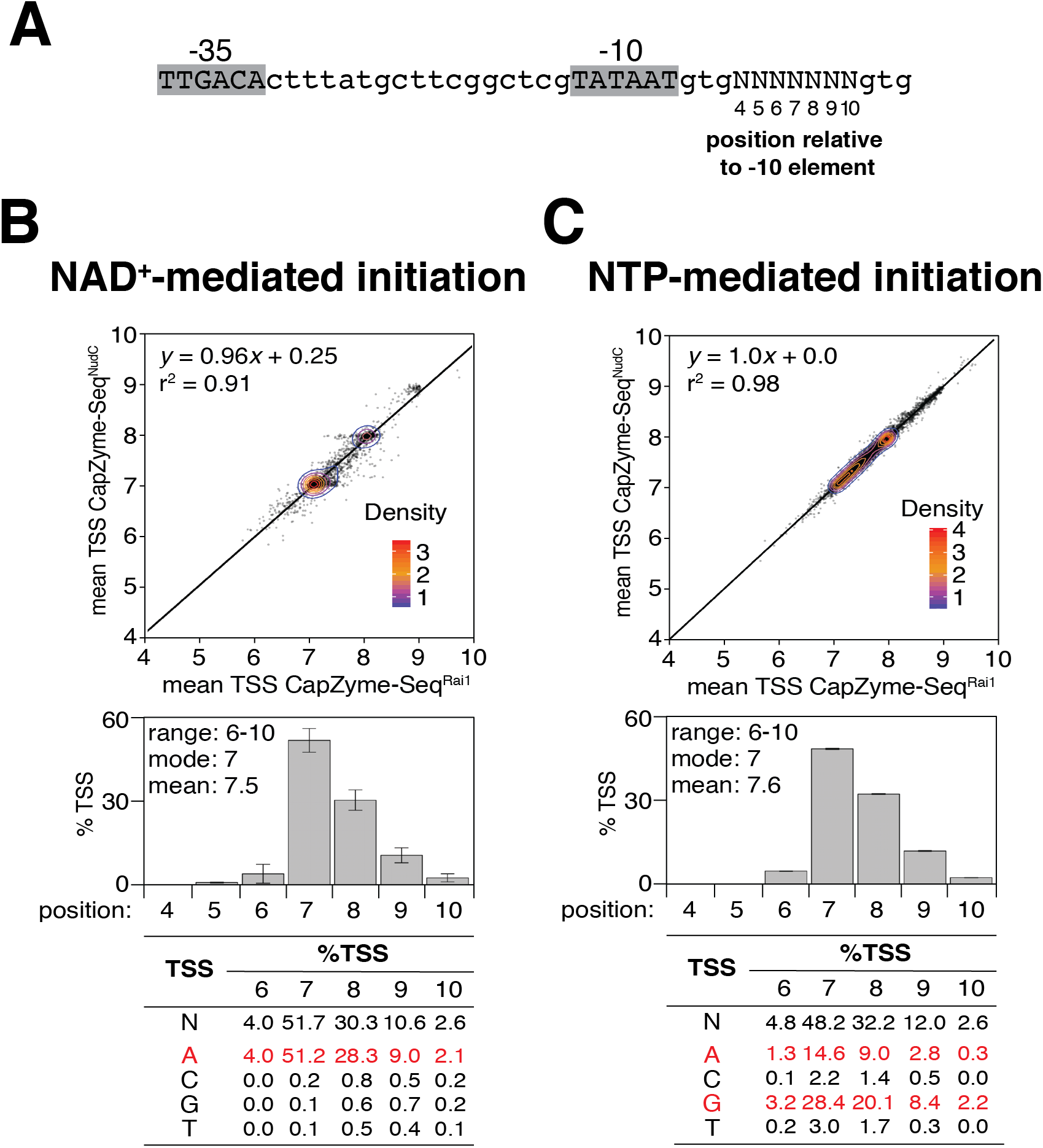
Determinants for transcription start site selection in NCIN capping with NAD^+^ *in vitro*. **A**. *lacCONS-N7* promoter library (4^7^, ~16,000 promoter sequences). **B-C**. Data for NAD^+^-mediated initiation (B) and NTP-mediated initiation (C). Top panels, mean TSS from CapZyme-Seq^NudC^ (n=3) vs. mean TSS from CapZyme-Seq^Rai1^ (n=3; mean TSS = [(4 x %TSS at position 4) + (5 x %TSS at position 5) +(6 x %TSS at position 6) + (7 x %TSS at position 7) + (8 x %TSS at position 8) + (9 x %TSS at position 9) + (10 x %TSS at position 10)] / 100). Middle panels, histograms of TSS positions (positions numbered relative to promoter -10 element; mean±SD of percentage of TSS at each position; n=6). Bottom panels, nucleotide frequencies for TSS selection at positions 6, 7, 8, 9, and 10 bp downstream of the −10 element (data for consensus nucleotides in red). See also Figure S1.

### Promoter sequence determinants for NCIN capping with NAD^+^ in vitro

The results in the previous section show that the modal, consensus TSS position for NAD^+^-mediated initiation is 7 bp downstream of the promoter −10 element and that the consensus TSS base pair for NAD^+^-mediated initiation is A:T. Considering the subset of ~4,000 promoter sequences in the *lacCONS-N7* promoter library that have A:T at the modal, consensus TSS position for NAD^+^-mediated initiation, 7 bp downstream of the promoter -10 element (A_+1_ promoters), we next assessed promoter sequence determinants for NAD^+^-mediated initiation at each of the three positions upstream of the TSS (positions 4, 5, and 6 bp downstream of the −10 element; positions −3, −2, and −1 relative to the TSS, position +1; Figure 4) and at each of the three positions downstream of the TSS (positions 8, 9, and 10 bp downstream of the −10 element; positions +2, +3, and +4 relative to the TSS; Figure 4). We find that capping efficiency depends on the identity of the nucleotide at each of these positions (Figures 4B,C and S2). At position −3, capping efficiency is higher for A, T, and C than for G, yielding the consensus H and anti-consensus G; at position −2, capping efficiency is higher for G and A than for T and C, yielding the consensus R and anti-consensus Y; at position −1, capping efficiency is higher for G and A than for T and C, yielding the consensus R; at position +2, capping efficiency is higher for G and C than for A and T, yielding the consensus S; at position +3, capping efficiency is higher for A and T than for G and C, yielding the consensus W and anti−consensus S; and at position +4, capping efficiency is higher for A and T than for G and C, yielding the consensus W and anti−consensus S (Figure 4C). The strongest dependence of capping efficiency on nucleotide identity, at positions other than the TSS, is observed at position -1. At this position, the mean relative capping efficiencies for promoters having A and G are ~2 to ~3 times higher than for promoters having T and ~7 to ~8 times higher than for promoters having C (Figure 4C). The second-strongest dependence of capping efficiency on nucleotide identity is observed at position +2. At this position, capping efficiencies for promoters having G and C are ~2 to ~3 times higher than for promoters having A and T (Figure 4C). The dependences of capping efficiencies on nucleotide identity at each of the other positions (−3, −2, +3 and +4) are smaller, but significant (Figure 4C).

**Figure 4.**
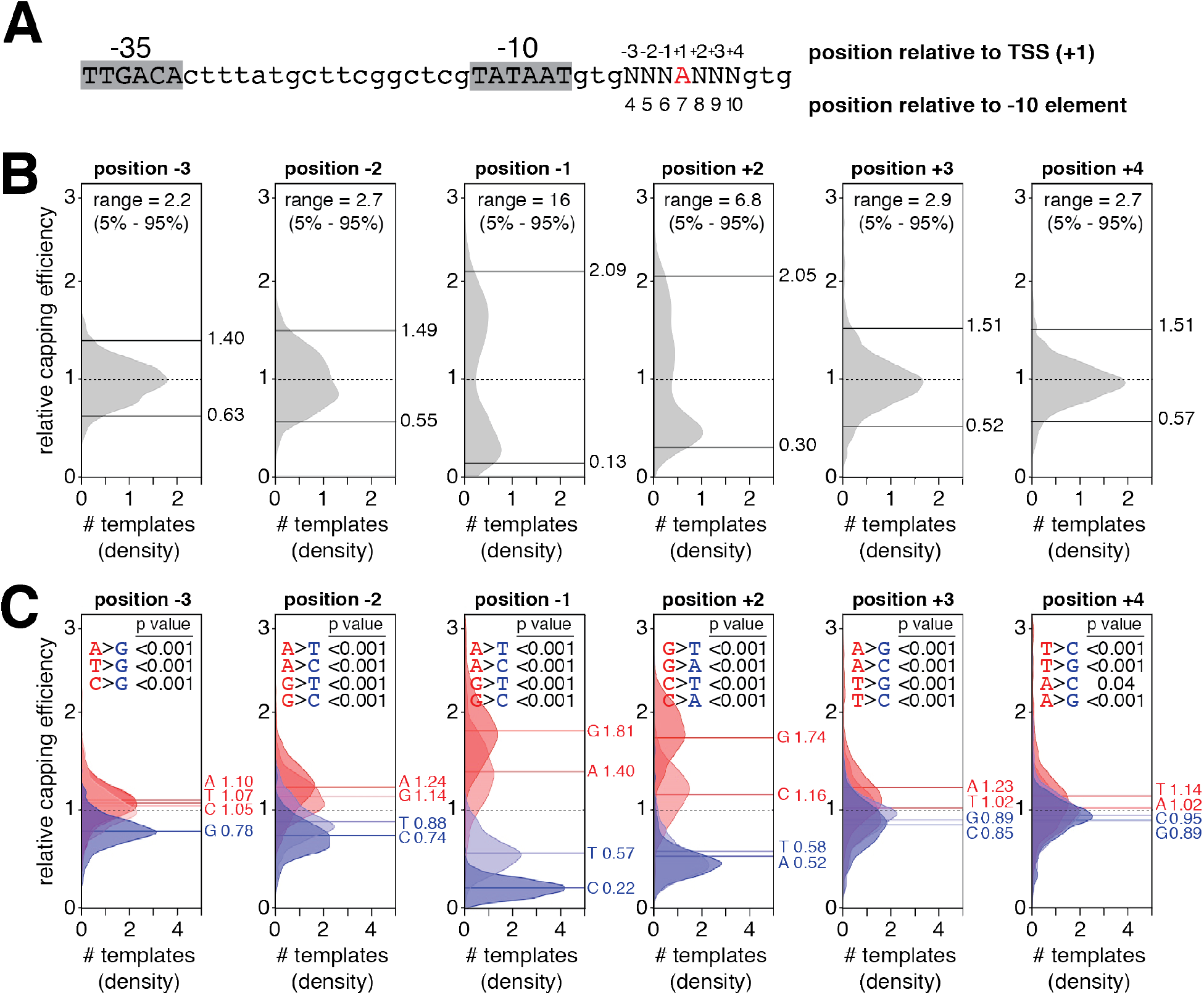
Promoter sequence determinants for NCIN capping with NAD^+^ *in vitro*. **A**. Subset of *lacCONS-N7* promoter library having A (red) at the position 7 bp downstream of -10 element (~4,000 sequences). **B**. Distributions of relative capping efficiency (calculated using the equation in Figure S2B) for ~4,000 A_+1_ promoter sequences at the positions immediately upstream of the TSS (positions -1, -2, and -3) and immediately downstream of the TSS (positions +2, +3, and +4). The dashed line is the mean relative capping efficiency, the upper and lower solid lines are the 95^th^ percentile and 5^th^ percentile, respectively, and the “range” is defined as the 95^th^ percentile relative capping efficiency divided by the 5^th^ percentile relative capping efficiency. **C**. Distributions of relative capping efficiency for ~4,000 A_+1_ promoter sequences parsed by position and nucleotide (A, T, C, or G). The dashed line is the mean relative capping efficiency for all sequences, the solid lines are the means for sequences having the indicated nucleotide. Distributions and lines are colored by consensus nucleotide (mean relative capping efficiency greater than 1; red) or anti-consensus nucleotide (mean relative capping efficiency less than 1; blue). Shown are p values for pairwise comparisons of consensus and anti-consensus nucleotides (Kolmogorov-Smirnov test). See also Figure S2.

To validate the CapZyme-Seq results, we analyzed capping efficiencies for individual promoters using a single-template gel assay [Figure S3; procedures as in (Bird et al., 2017)]. The results of the ~1,000-template CapZyme-Seq assays and the single-template gel assays were in full agreement (Figure S3).

The finding that position −1 is a crucial sequence determinant, with G or A as the preferred nucleotides, confirms our previous results (Bird et al., 2016) and contradicts results of (Julius and Yuzenkova, 2017). Julius and Yuzenkova did not observe specificity at position −1, most likely because they measured only 1/K_m_, and not k_cat_/K_m_, and thus were unable to detect specificity manifest at the level of k_cat_. The findings for sequence specificity at positions −3, −2, +2, +3, and +4 are new to this work.

### Promoter consensus sequence for NCIN capping with NAD^+^ in vitro

The results in the previous section provide a promoter consensus and anti-consensus sequence for NCIN capping with NAD^+^: H_−3_R_−2_R_−1_;A_+1_S_+2_W_+3_W_+4_ and G_−3_Y_−2_Y_−1_A_+1_W_+2_S_+3_S_+4_, respectively (Figure 5A). The results in Figure 5B indicate that the mean relative capping efficiencies for promoters having a consensus nucleotide at positions −3, −2, −1, +2, +3 and +4 are ~1.4-fold, ~1.5-fold, ~4.1-fold, ~2.6-fold, ~1.3-fold, and ~1.2-fold, respectively, greater than the mean relative capping efficiencies for promoters having an anti-consensus nucleotide at these positions. Mean capping efficiency values for consensus A+1 promoter sequences vs. anti-consensus A+_1_ promoter sequences differ by ~40-fold (Figure 5C). Single-template gel assays comparing a consensus A+1 promoter and anti-consensus A+1 promoter confirm the ~40-fold difference in capping efficiency observed by CapZyme-Seq (Figure 5D).

**Figure 5.**
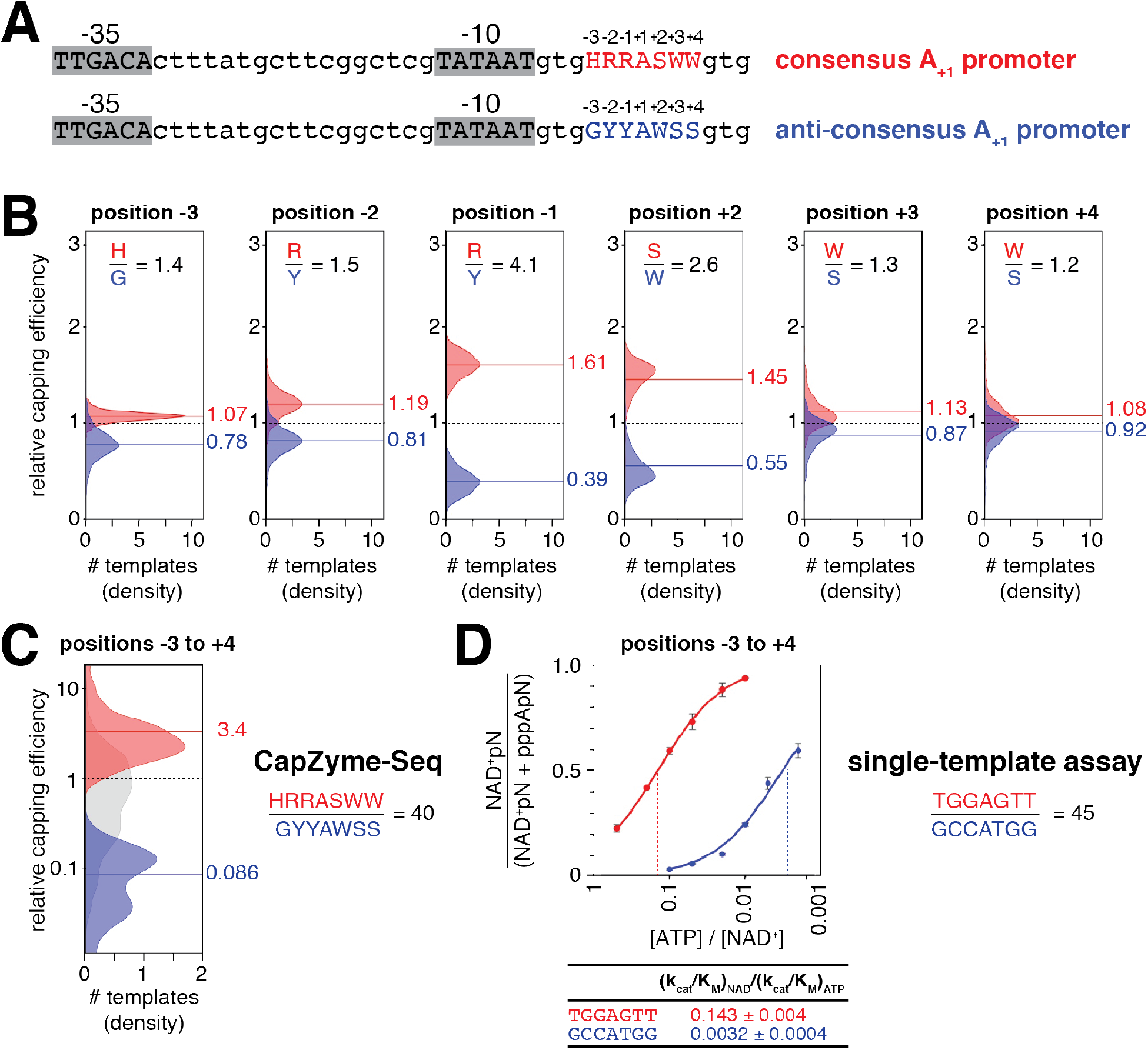
Promoter consensus sequence for NCIN capping with NAD^+^ *in vitro.* **A**. *lacCONS* promoter derivatives with consensus A_+1_ sequence (red) and anti-consensus A_+1_ sequence (blue) for NAD^+^ capping. **B**. Distributions of relative capping efficiency, for ~4,000 A_+1_ promoter sequences parsed by position and nucleotide (H, G, R, Y, S, or W) and colored by consensus nucleotide (red) or anti-consensus nucleotide (blue). The dashed line is the mean relative capping efficiency for all sequences, the solid lines are the means for sequences having a consensus nucleotide (red) or an anti-consensus nucleotide (blue). **C**. Distributions of relative capping efficiency for consensus A_+1_ sequences (red), anti-consensus A_+1_ sequences (blue), or all A_+1_ sequences (grey). **D**. Dependence of NAD^+^ capping on [ATP] / [NAD^+^] ratio for representative consensus A_+1_ promoter sequence (red) and anti-consensus A_+1_ promoter sequence (blue): data from single-template gel assays (mean±SD; n=3). See also Figure S3.

### Strand specificity of NCIN capping with NAD^+^ in vitro

In a catalytically competent transcription initiation complex (RNAP-promoter open complex), positions −3, −2, −1, +1, and +2 are part of an unwound, non-base-paired “transcription bubble” (Zhang et al., 2012). This raises the question whether specificity for positions −3, −2, −1, +1, and +2 is carried by the single-stranded “template strand” of the transcription bubble (the strand that templates incoming nucleotide substrates), the single stranded “non-template strand” of the transcription bubble, or both (Figure 6A). To address this question for the TSS, position +1, we analyzed NAD^+^ capping for promoter derivatives having consensus nucleotides at this position on both the template and non-template strands, on the template strand only, on the non-template strand only, or on neither (Figure 6B). NAD^+^ capping was observed when consensus nucleotides were present on both strands or on the template strand only, but not when consensus nucleotides were present on the non-template strand only or on neither strand (Figure 6B). We conclude that, at position +1, the sequence information for NAD^+^ capping resides exclusively in the template strand. To address this question for positions −3, −2, −1, and +2, we compared NAD^+^ capping efficiencies for heteroduplex promoter derivatives having the consensus or anti-consensus nucleotides on the template strand and an abasic site (*) on the non−template strand (Figure 6C). For each position, the capping efficiency ratio for constructs having the consensus vs. anti-consensus only on the template strand matched the capping efficiency ratio for homoduplex promoter derivatives having consensus vs. anti-consensus on both strands (Figure 6C). We conclude that, as at position +1, at positions −3, −2, −1, and +2, the sequence information for NAD^+^ capping resides exclusively in the template strand.

**Figure 6.**
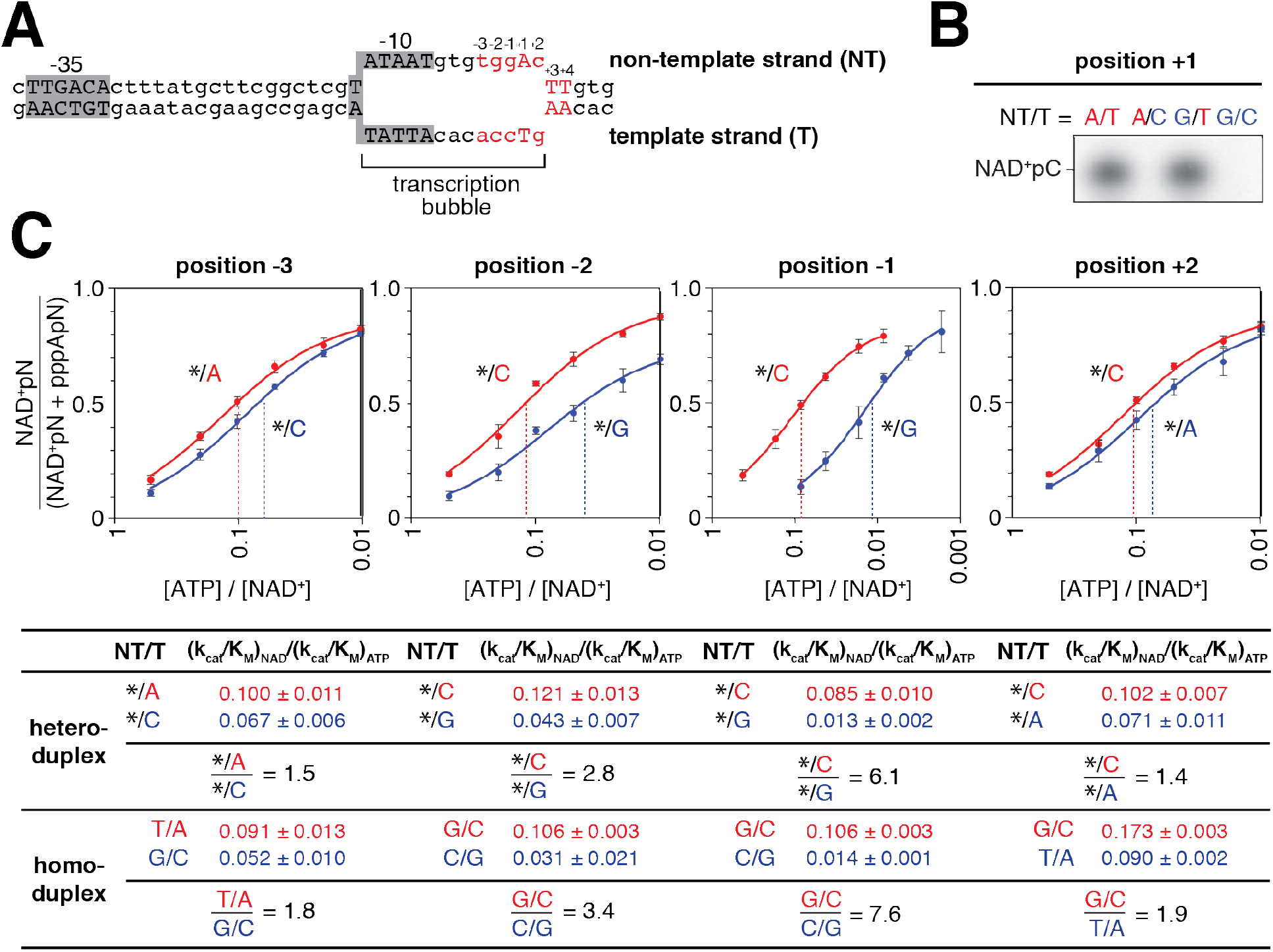
Strand specificity of NCIN capping with NAD^+^ *in vitro*. **A**. *lacCONS* promoter derivative containing consensus A_+1_ promoter sequence in context of RNAP-promoter open complex. DNA non-template strand (NT) on top; DNA template strand (T) on bottom; Unwound, non-base-paired DNA region, “transcription bubble,” indicated by raised and lowered nucleotides **B**. Products of transcription reactions with NAD^+^ as initiating nucleotide and [α^32^P]-CTP as extending nucleotide for templates having the consensus nucleotides at the TSS, position +1, on both DNA strands, non-template strand only, template strand only, or neither. **C**. Dependence of NAD^+^ capping on [ATP] / [NAD^+^] ratio for templates having an abasic site (*) on the DNA non-template strand and either consensus base (red) or anti-consensus base (blue) on the DNA template strand at each of positions -3, -2, -1, and +2, relative to TSS. Below, capping efficiencies and consensus/anti-consensus capping efficiency ratios for heteroduplex templates with an abasic site on the DNA non-template strand or, for comparison, for homoduplex templates having a complementary nucleotide on the DNA non-template strand (mean±SD; n=3). See also Figure S5.

### CapZyme-Seq analysis of NCIN capping in vivo

To demonstrate the utility of CapZyme-Seq for analysis of RNA isolated from living cells we applied CapZyme-Seq to (i) define promoter sequence determinants for NCIN capping *in vivo* in *E. coli*, and (ii) to identify and quantify small RNAs (sRNAs) *in vivo*. Both of these objectives were pursued, in parallel, using RNA isolated from a single culture.

### Promoter-sequence determinants for NCIN capping in vivo

To determine total levels of NCIN capping and to define promoter-sequence determinants for NCIN capping we isolated RNA products from *E. coli* cells containing the MASTER *lacCONS-N7* template library used in the experiments in Figures 2–5 and analyzed RNA products from the 4^7^ MASTER *lacCONS-N7* promoter sequences using CapZyme-Seq^NudC^ (Figure 7A). We observed NCIN capping with many promoter sequences *in vivo*, extending and generalizing our conclusion from previous work with a single promoter sequence (Bird et al. 2016). The level of NCIN capping differs for RNA products from the 4^7^ different promoter sequences, ranging from 0 to 38%, with a mean of 3%, and a median of 2% (Figure 7A). We see a broad distribution of percent capping values *in vivo* (Figure 7A) reminiscent of the broad distribution of percent capping values observed *in vitro* (Figure 2C).

**Figure 7.**
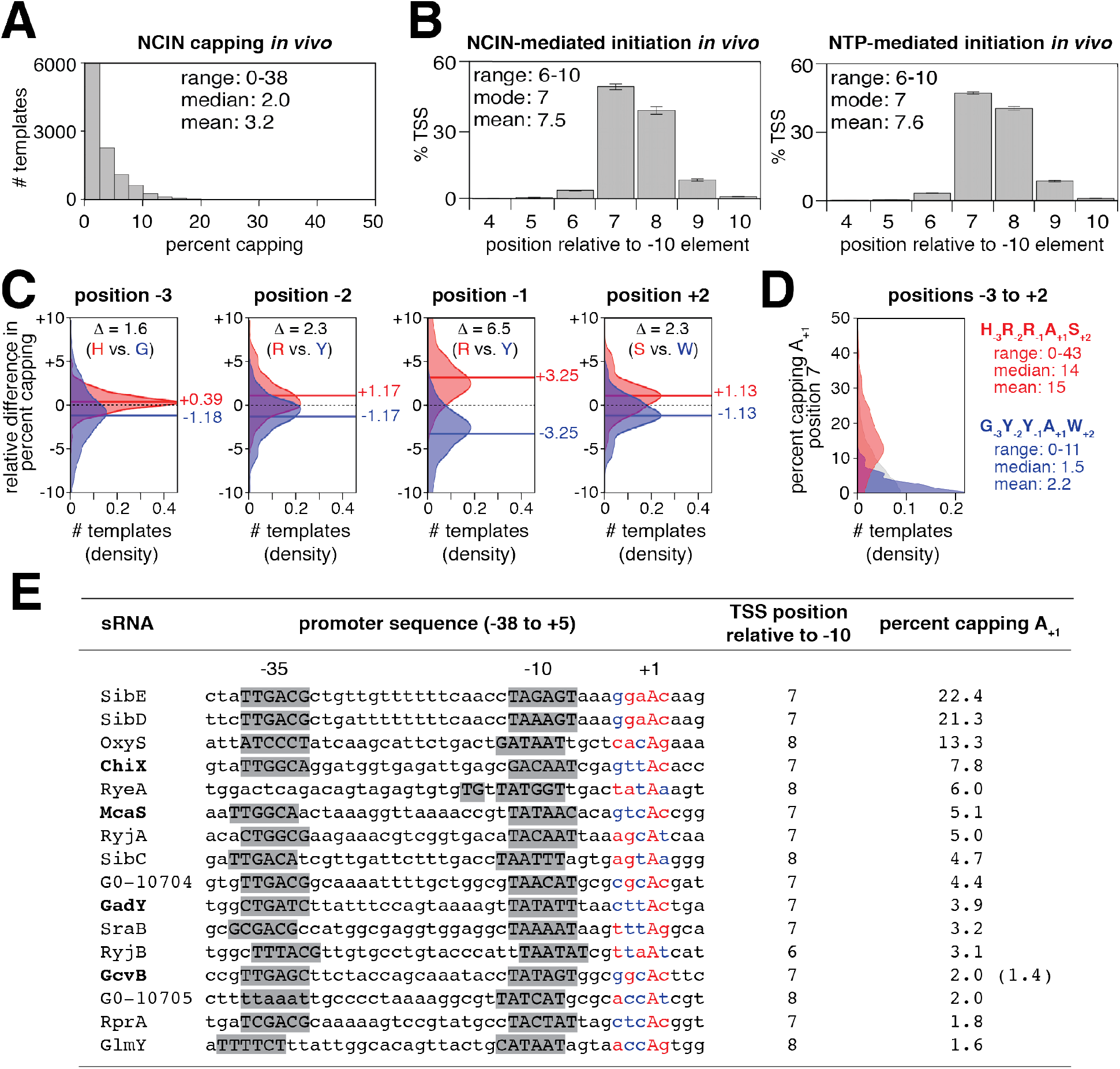
CapZyme-Seq analysis of NCIN capping *in vivo*. **A-D**. Promoter sequence determinants for NCIN capping *in vivo* in *E. coli*. Percent capping histograms (A); TSS position histograms (B; mean±SD of percentage of TSS at each position; n=3); relative percent capping difference distributions (C; calculated using the equation in Figure S4B; the dashed line is 0, the solid lines are the means, consensus nucleotides are colored red, anti-consensus nucleotides are colored blue); percent capping histograms for −3 through +2 consensus (red) and anti-consensus (blue) sequences for NCIN-capping *in vivo* (D). **E**. Identification and quantitation of NCIN-capped sRNAs *in vivo* in *E. coli*. Bold indicates the four sRNA sequences previously identified as NAD^+^ capped (Cahova et al., 2015). In the promoter sequences, grey shading represents the promoter -10, extended -10, and -35 promoter elements, and colors indicate matches to the -3 through +2 consensus (red) and anti-consensus (blue) sequences for NCIN capping *in vivo*. Percent capping values represent the mean of three independent measurements. In the column for percent capping the number reported previously in (Nubel et al., 2017) for GcvB is in parentheses. See also Figure S4.

The preferred TSS positions for NCIN capping *in vivo* matches the preferred TSS positions for NCIN capping with NAD^+^ *in vitro* (mode = 7 bp downstream of −10 element; mean = 7.5 bp downstream of −10 element; Figures 3B and 7B). The preferred TSS base pair for NCIN capping *in vivo* (A:T) also matches the preferred TSS base pair for NCIN capping with NAD^+^ *in vitro* (Figures 3B and S4A).

Considering the subset of ~4,000 promoter sequences in the *lacCONS-N7* promoter library that have A:T at the modal, consensus TSS position for NCIN capping, 7 bp downstream of the promoter −10 element (A_+1_ promoters), we next assessed promoter sequence determinants for NCIN capping *in vivo* at each of the three positions upstream and downstream of the TSS (Figures 7C and S4C-D). We find that, for positions −3 to +2, the sequence determinants for NCIN capping *in vivo* (Figures 7C and S4C) match those for NCIN capping with NAD^+^ *in vitro* (Figures 4, 5, and S3). At position +3 we observe no sequence preferences *in vivo* (Figure S4D) and at position +4 we observe sequence preferences similar to, but weaker than those observed *in vitro* (Figures 4 and S4D). The results provide promoter consensus and anti-consensus sequences for NCIN capping *in vivo*―H_−3_R_−2_R_−1_A_+1_S_+2_ and G_−3_Y_−2_Y_−1_A_+1_W_+2_ (Figure 7C,D)—that match the corresponding promoter consensus and anti-consensus sequences for positions −3 to +2 for NCIN capping with NAD^+^ *in vitro* (Figure 5). The strongest promoter sequence dependence of NCIN capping *in vivo*, apart from that at the TSS, is observed at position −1 (Figure 7C), matching the pattern observed for NCIN capping with NAD^+^ *in vitro* (Figure 4B,C). At this position, the difference, Δ, in mean percent capping for consensus vs. anti-consensus is ~6.5% (Figure 7C). At positions −3, −2, and +2, the difference, Δ, in mean percent capping for consensus vs. anti-consensus is ~2%. Considering positions −3 through +2, the difference, Δ, in mean percent capping for consensus vs. anti-consensus is ~13% (Figure 7D).

### Identification and quantitation of NCIN-capped small RNAs in vivo

Identities of several NAD^+^-capped small RNA (sRNA) and sRNA-like 5′-RNA fragments in *E. coli* total cellular RNA have been reported (Cahova et al., 2015). Here, we applied CapZyme-Seq to identify and quantify NCIN capping of all annotated *E. coli* sRNAs. We isolated *E. coli* total cellular RNA and performed CapZyme-Seq^NudC^ using primers for the cDNA synthesis step designed to target 77 annotated sRNAs of *E. coli* (Keseler et al., 2017). Analysis of uncapped, 5’-triphosphate RNA using Rpp identified 16 sRNAs arising from promoters having A:T at the TSS position (Figure 7E). Analysis of NCIN-capped RNA using NudC shows that all 16 sRNAs exhibit NCIN capping, including the four native *E. coli* sRNAs previously identified as NCIN capped (ChiX, McaS, GadY, and GcvB; Figure 7E). NCIN capping levels for the 16 sRNAs ranged from 1.6% to 22.4%. Three RNAs shows particularly high NCIN capping levels: SibE, SibD and OxyS (22.4%, 21.3%, and 13.3%). SibE and SibD are anti-toxin sRNAs and OxyS is an sRNA regulator of oxidative stress. All three most highly NCIN-capped sRNAs are newly identified in this work. We note that the two most highly capped sRNAs are transcribed from promoters that match the four most important positions, −2 to +2, of our consensus sequence for NCIN capping *in vivo*.

### Proposed basis for promoter sequence specificity for NCIN capping with NAD^+^

The results in Figures 3–6 show that the efficiency of NCIN capping with NAD^+^ is determined by sequence at the TSS (+1), sequence at the three positions immediately upstream of the TSS (−3 to −1), and sequence at the three positions immediately downstream of the TSS (+2 to +4). There is an essentially absolute preference for A at the TSS (Figure 3B). At the first position upstream of the TSS, position −1, sequence has very strong effects on efficiency of NAD^+^ capping (up to at least 16-fold; Figure 4B). At the second and third positions upstream of the TSS, positions −2 and −3, sequence has small, but significant, effects (up to at least 2.7-fold and 2.2-fold, respectively; Figure 4B). At the first position downstream of the TSS, position +2, sequence has large effects (up to at least 6.8-fold; Figure 4B). At the next two positions downstream of the TSS, positions +3 and +4, promoter sequence has small but significant effects (up to at least 2.9-fold and 2.7-fold, respectively; Figure 4B). The results in Figure 6 show that the specificity determinants at positions −3 to +2 reside exclusively in the template strand of the promoter DNA within the unwound transcription bubble of the RNAP-promoter open complex.

The essentially absolute preference for an A:T base pair at the at the TSS, position +1, results from the Watson-Crick base-paring preference of the adenosine moiety of NAD^+^ with a T on the template-strand (Figure 1A, and Figure S5A).

Structural modeling suggests that the very strong preference for R (Y on template strand) at the position immediately upstream of the TSS, position −1, can be understood in terms of “pseudo-base pairing” of the NAD^+^ nicotinamide moiety with the DNA template strand base at position −1 (Figure S5). The NAD^+^ nicotinamide can be positioned to form a nicotinamide:Y pseudo-base pair with template-strand C at position −1 or, with a 180° rotation about the pyridine-amide bond of the NAD^+^ nicotinamide, with template-strand T at position −1, in each case, forming two H-bonds with Watson-Crick H-bonding atoms of the template-strand and stacking on the NAD^+^ adenine base (Figure S5A,B). In contrast, the NAD^+^ nicotinamide moiety would experience severe steric clash with template strand A or G at position −1 (Figure S5C).

Structural modeling suggests that the specificity for R (Y on template strand) at position −2 also can be understood in terms of stacking preferences for pseudo-base pairing by the NAD^+^ nicotinamide moiety to the template-strand base at position −1. A template-strand Y at position −2 can stack favorably on the NAD^+^ nicotinamide moiety of a nicotinamide:Y pseudo-base pair (Figure S5A), but a template-strand R at position −2 would clash with the NAD^+^ nicotinamide moiety.

The strong specificity for S (S on template strand) at the first position downstream of the TSS, position +2, can be understood in terms of differences of 1/K_m_ for the incoming extending NTP, which base pairs with the template-strand base at position +2 (Rhodes and Chamberlin, 1974; Jensen et al., 1986), together with different sensitivities to this parameter of NAD^+^-mediated initiation to ATP-mediated initiation.

The specificity for W:W base pairs at positions +3 and +4 observed *in vitro*, potentially can be understood in terms of differences in DNA duplex stabilities, and corresponding DNA unwinding energies for W:W base pairs vs. S:S base pairs, together with different sensitivities to these parameters of NAD^+^-mediated initiation to ATP-mediated initiation.

### Prospect

Jaschke and co-workers have reported a method that combines click-chemistry covalent capture with high-throughput sequencing to isolate and identify NAD^+^-capped RNAs, “NAD^+^-capture-seq” (Cahova et al., 2015; Winz et al., 2017). NAD^+^-capture-seq has enabled detection of NAD^+^ capped RNAs in bacteria, yeast, and human cells in culture (Cahova et al., 2015; Jiao et al., 2017; Walters et al., 2017). However, NAD^+^-capture-seq does not enable single-nucleotide resolution identification of RNA 5′-ends, does not enable quantitation of relative yields of NAD^+^-capped and uncapped RNA, and does not enable detection of RNAs carrying NCIN caps other than NAD^+^. In contrast, the method we report in this work, “CapZyme-Seq” (Figure 1), which combines selective enzymatic processing of NCIN-capped 5′ ends and uncapped 5′ ends with high-throughput sequencing, enables single-nucleotide resolution identification of RNA 5′-ends (Figures 3, S1, 7B and S4A), enables quantitation of relative yields of NAD^+^-capped and uncapped RNA (Figures 2, 4, 5, and 7), and enables detection of RNAs carrying NCIN-caps other than NAD^+^ (Figure 1).

By combining CapZyme-Seq with multiplexed transcriptomics, we determined the efficiencies of NAD^+^ capping by *E. coli* RNAP σ^70^ holoenzyme for ~16,000 promoter sequences (Figure 2). A priority for future studies will be to adapt the methods employed in this work to define the promoter-sequence determinants for NAD^+^ capping by bacterial RNAP holoenzymes carrying alternative σ factors, archaeal RNAP, eukaryotic RNAP I, II, and III, mitochondrial RNAP, and chloroplast RNAP.

The prevalence of nucleoside-containing metabolites that can function as NCINs underscores the need to determine, for each NCIN, the promoter-sequence determinants for NCIN capping. The method reported here, either using the same decapping enzymes or using alternative decapping enzymes with alternative decapping specificities, should enable analysis of determination of NCIN-capping efficiencies and promoter sequence determinants thereof for each nucleoside-containing metabolite that can serve as an NCIN (e.g., NADH, dpCoA, FAD, UDP-glucose, and UDP-GlcNAc).

NCIN caps provide a layer of epitranscriptomic regulation by modulating RNA stability and translation efficiency. Understanding the full impact of NCIN capping as a mechanism for altering RNA fate requires understanding the mechanism(s) by which the distributions of NCIN caps for different RNA products are determined—i.e. the mechanism(s) of “NCIN targeting.” The method reported here was developed, validated, and applied to RNA generated *in vitro* and *in vivo* in *E. coli*. This same method should be applicable, essentially without modification, to RNA isolated from any source, including eukaryotic cells, tissues, organs, and organisms.

## ACKNOWLEDGEMENTS

Work was supported by Czech Science Foundation grant 15-05228S (L.K.), and by NIH grants GM067005 (M.K), GM041376 (R.H.E), and GM118059 (B.E.N).

## AUTHOR CONTRIBUTIONS

Conceptualization, M.K., R.H.E, and B.E.N.; Methodology, Yu Zhang, I.B., L.K., and B.E.N.; Formal Analysis, J.B., Yuanchao Zhang, M.K., D.M.T., R.H.E. and B.E.N.; Investigation, J.G.B., I.O.V., and X.J.; Data Curation, Yuanchao Zhang; Writing - Original Draft, R.H.E. and B.E.N.; Writing - Review & Editing, J.G.B., I.O.V., L.K., and M.K.; Visualization, J.G.B., Yuanchao Zhang, and B.E.N.; Supervision, L.K., M.K., D.M.T., R.H.E., and B.E.N.; Project Administration, R.H.E. and B.E.N.; Funding Acquisition, L.K., M.K., R.H.E., and B.E.N. The authors declare that no competing interests exist.

---

**Figure S1 (Related to Figure 3).**
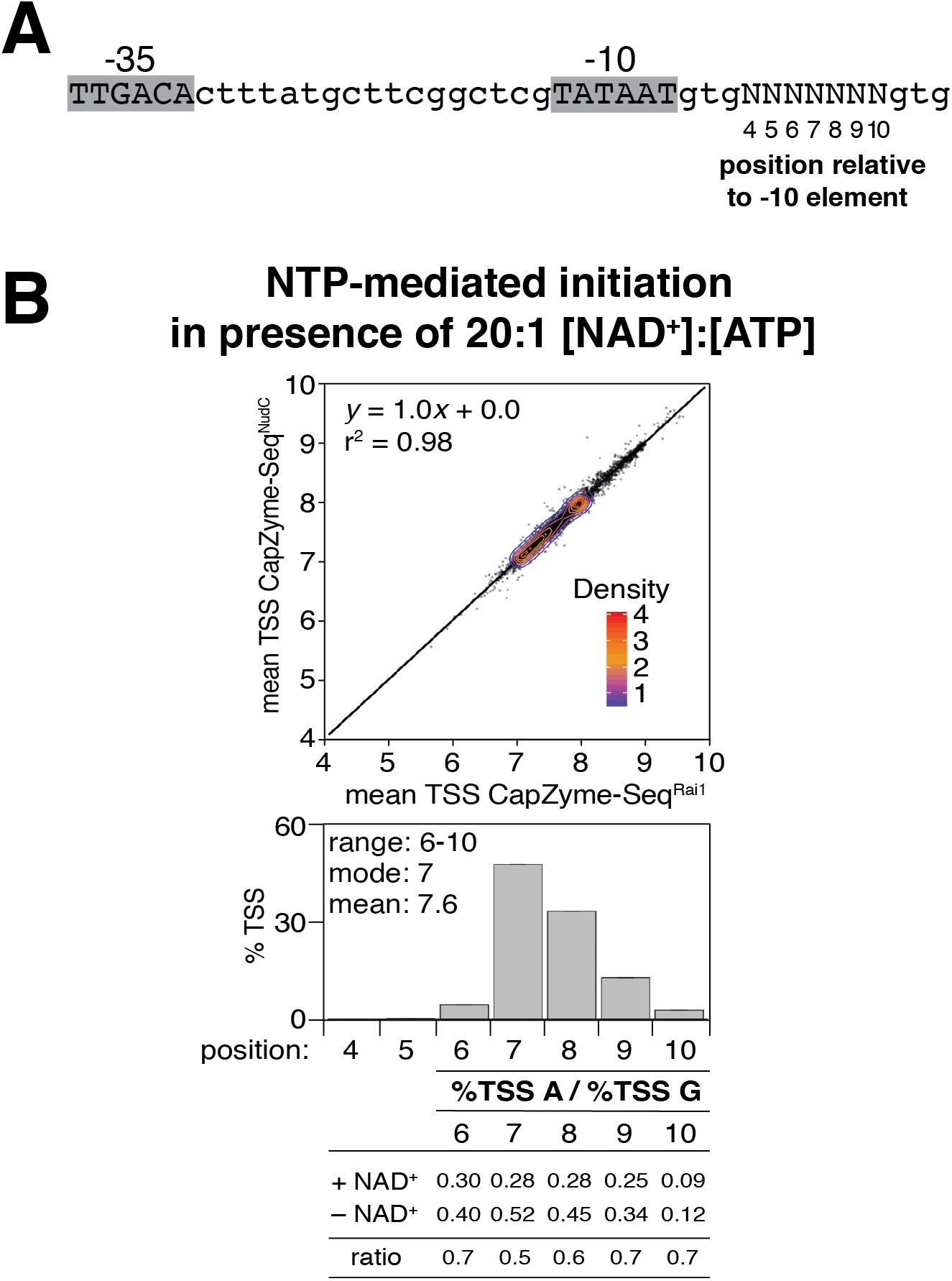
Transcription start site selection in NTP-mediated initiation in the presence of NAD^+^ *in vitro*. **A**. *lacCONS-N7* promoter library (4^7^, ~16,000 promoter sequences). **B**. Data for NTP-mediated initiation in the presence of NAD^+^. Top panel, mean TSS from CapZyme-Seq^NudC^ (n=3) vs. mean TSS from CapZyme-Seq^Rai1^ (n=3; mean TSS = [(4 x %TSS at position 4) + (5 x %TSS at position 5) +(6 x %TSS at position 6) + (7 x %TSS at position 7) + (8 x %TSS at position 8) + (9 x %TSS at position 9) + (10 x %TSS at position 10)] / 100). Middle panel, histograms of TSS positions (positions numbered relative to promoter −10 element; mean±SD of percentage of TSS at each position; n=6). Bottom panel, nucleotide frequencies for TSS selection with A vs. G in the absence or presence of NAD^+^ and ratios of nucleotide frequencies for TSS selection with A vs. G in the presence or absence of NAD^+^.

**Figure S2 (Related to Figure 4).**
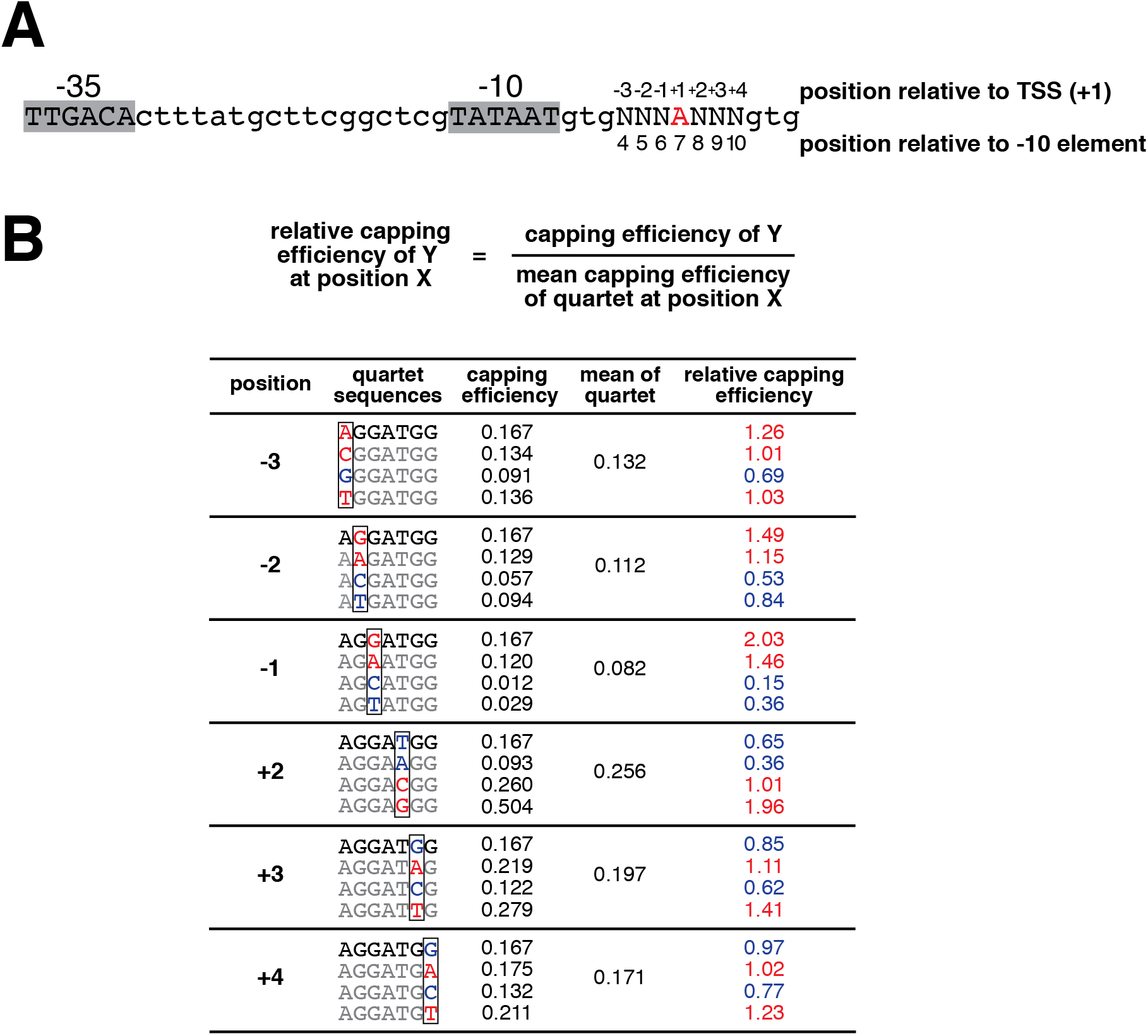
Promoter sequences determinants for NCIN capping with NAD^+^: determination of relative capping efficiency *in vitro*. **A**. Subset of *lacCONS-N7* promoter library having A (red) at the position 7 bp downstream of -10 element (A_+1_ promoter sequences; ~4,000 sequences). **B**. Determination of relative capping efficiencies at positions -3 through +4 for representative A_+1_ promoter sequences. Relative capping efficiency at each position was determined for groups of four promoter sequences, “quartets,” having A, G, C, or T, along with sequences identical at each other position. The relative capping efficiency of each A_+1_ promoter sequence, Y, at each position, X, is calculated by dividing the capping efficiency of Y by the mean capping efficiency of the quartet that Y belongs to at position X. Table shows the determination of relative capping efficiencies at positions −3 through +4 for a representative A_+1_ sequence (AGGATGG; black). Listed in grey are quartet sequences other than AGGATGG for each position. Also shown are the mean capping efficiency of each sequence (n=6), the mean capping efficiency of each quartet; and the relative capping efficiency calculated using the equation above the Table. The position that varies for each quartet is boxed and the nucleotide (A, T, C, or G) colored by consensus nucleotide (relative capping efficiency greater than 1; red) or anti-consensus nucleotide (mean relative capping efficiency less than 1; blue).

**Figure S3 (Related to Figure 5).**
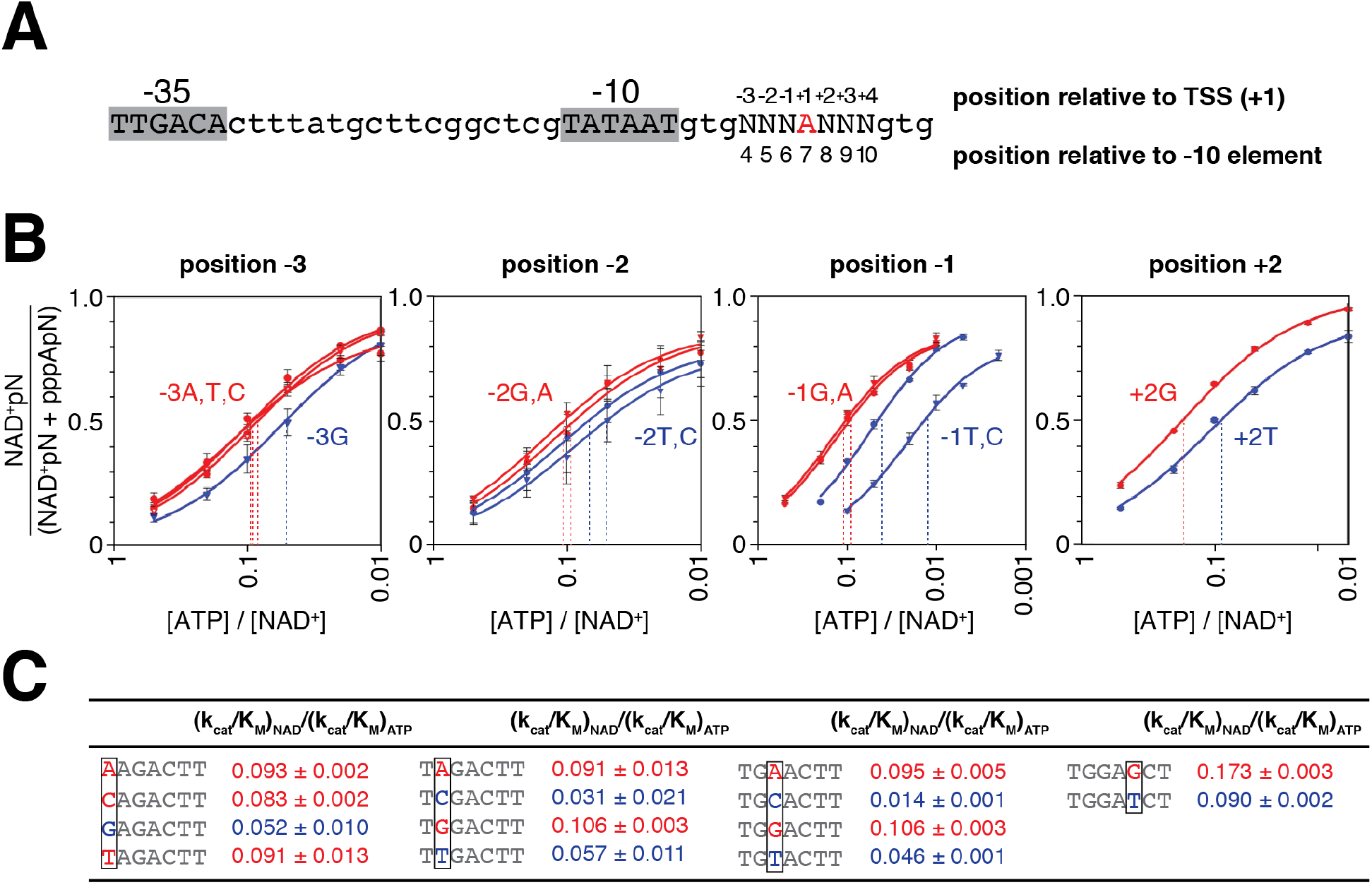
Promoter sequence determinants for NCIN capping with NAD^+^ *in vitro*: single-template gel data. **A**. Subset of *lacCONS-N7* promoter library having A (red) at the position 7 bp downstream of −10 element (~4,000 sequences). **B**. Dependence of NAD^+^ capping on [NAD^+^] / [ATP] ratio determined in single-gel assays: effects of promoter sequence at positions −3, −2, −1 and +2 relative to TSS. The nucleotide (A, T, C, or G) and the best fit line is colored by consensus nucleotide (red) or anti-consensus nucleotide (blue). **C**. Capping efficiencies from single-template gel assays (mean±SD; n=3). The position that varies for each set of sequences is boxed. The nucleotide (A, T, C, or G) and capping efficiency value for each sequence is colored by consensus nucleotide (red) or anti-consensus nucleotide (blue).

**Figure S4 (Related to Figure 7).**
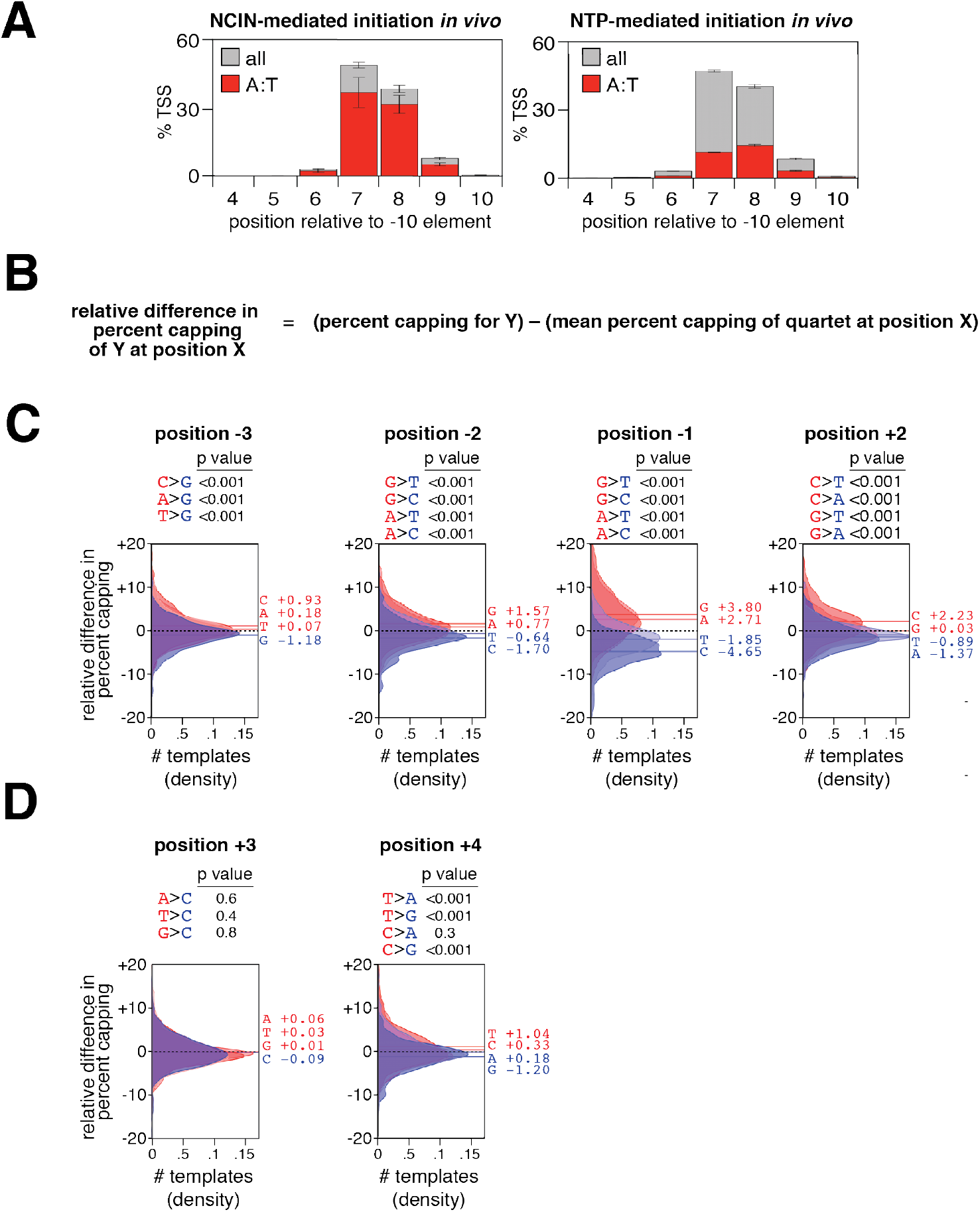
Promoter sequence determinants for NCIN capping *in vivo* in *E. coli*. **A**. TSS position histograms (mean±SD of percentage of TSS at each position; n=3; red, TSS is an A:T base pair; grey all TSS). **B**. Equation for calculating relative difference in percent capping. **C**. Percent capping difference distributions for positions −3 through +2. Distributions of relative percent capping differences for ~4,000 A_+1_ promoter sequences parsed by position and nucleotide (A, T, C, or G). The dashed line is at 0, the solid lines are the mean relative percent capping differences for sequences having the indicated nucleotide. Distributions and lines are colored by consensus nucleotide (mean relative percent capping difference of greater than 0; red) or anti-consensus nucleotide (mean relative percent capping difference of less than 0; blue). Shown are p values for pairwise comparisons of consensus and anti-consensus nucleotides (Kolmogorov-Smirnov test). **D**. Percent capping difference distributions for positions +3 and +4. Distributions and lines are colored by nucleotides with mean relative percent capping differences of greater than 0 (red) or less than 0 (blue). Shown are p values for pairwise comparisons of the indicated nucleotides (Kolmogorov-Smirnov test).

**Figure S5 (Related to Figure 6).**
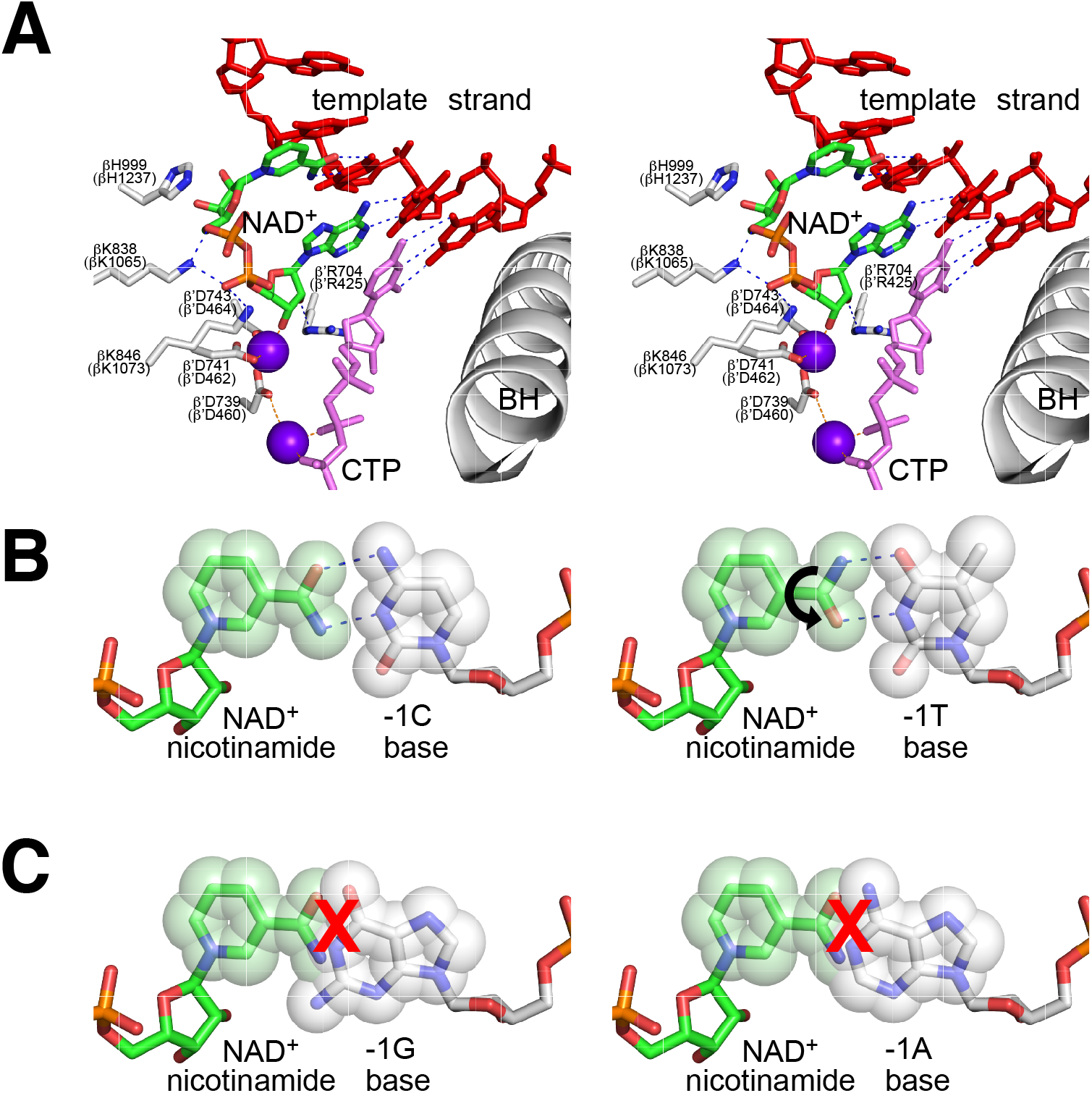
Proposed basis for promoter-sequence specificity for NCIN capping with NAD^+^: positions +1, −1 and −2. **A**. Stereoview of structural model of substrate complex for NAD^+^-mediated initiation, showing base pairing of NAD^+^ adenine moiety with DNA template-strand base T at position +1, pseudo-base pairing by NAD^+^ nicotinamide moiety with DNA template-strand base C at position −1, and stacking of NAD^+^ nicotinamide moiety with DNA template strand base C at position −2. NAD^+^ in green, blue, red, and orange; extending nucleotide CTP in pink; DNA template strand in red; RNAP active-center residues that contact NAD in grey, blue, and red; RNAP active-center catalytic Mg^2+^(I) and Mg^2+^(II) as violet spheres; and RNAP active-center bridge helix as grey ribbon labeled BH. Dashed lines, H-bonds. **B**. Structural model of pseudo-base pair between NAD^+^ nicotinamide and pyrimidine base C at template strand-position −1 or, with 180° rotation (black curved arrow) about pyridine-amide bond of NAD nicotinamide moiety, with pyrimidine base T at template strand-position −1. **C**. Structural model of steric clash (red X) between NAD^+^ nicotinamide moiety and purine base G or A at template strand position −1.

## METHODS

### CONTACT FOR REAGENT AND RESOURCE SHARING

Requests for further information or reagents should be directed to Bryce E. Nickels.

## METHOD DETAILS

### Proteins

*E. coli* RNAP core enzyme used in transcription experiments was prepared from *E. coli* strain NiCo21(DE3) transformed with plasmid pIA900 (Svetlov and Artsimovitch, 2015) as described (Artsimovitch et al., 2003). *E. coli* σ^70^ used in transcription experiments was prepared from *E. coli* strain BL21(DE3) transformed with plasmid pσ^70^-His (gift of J. Roberts) as described (Marr and Roberts, 1997). RNAP holoenzyme was formed by mixing 2 μM RNAP core and 10 μM σ^70^ in 10 mM Tris-Cl, pH 8.0, 100 mM KCl, 10 mM MgCl_2_, 0.1 mM EDTA, 1 mM DTT, and 50% glycerol and incubating for 30 min at 25°C.

*E. coli* NudC was prepared from *E. coli* strain NiCo21(DE3) transformed with plasmid pET NudC-His (Bird et al., 2016) as described (Cahova et al., 2015). *S. pombe* Rai1 was prepared from *E. coli* strain BL21(DE3) Rosetta as described (Xiang et al., 2009). 5′ RNA polyphosphatase (Rpp) was purchased from Epicentre.

### Oligonucleotides

A complete list of oligodeoxyribonucleotides and oligoribonucleotides are provided in the Key Resources Table. All oligodeoxyribonucleotides and oligoribonucleotides were purified with standard desalting purification. Adapters and primers used in MASTER library construction were also HPLC purified.

Linear *in vitro* transcription templates for single-template gel assays were generated by PCR using Phusion HF Polymerase master mix 5 nM of template oligo, 0.5 mM of forward primer and 0.5 mM of reverse primer. PCR products were purified using a QIAquick PCR purification kit prior to use in transcription reactions.

Templates with abasic sites on the DNA non-template strand (Figure 6) were generated by mixing 1.1 μM non-template strand oligo with 1 μM template strand oligo in 10 mM Tris pH 8.0. Mixtures were heated to 95^°^C for 10 minutes, cooled to 25^°^C, and incubated for 20 minutes using a Dyad PCR machine (Bio-Rad).

### NudC, Rai1, and Rpp processing assays

For assays shown in Figure 1C, radiolabeled NCIN-capped RNA products were generated by *in vitro* transcription. 10 nM linear *lacCONS* A-less cassette template (made by PCR using oligos JB277, JB285 and JB455) was mixed with 50 nM RNAP holoenzyme in 10 mM Tris HCl pH 8.0, 50 mM NaCl, 10 mM MgCl_2_, 0.1 mM EDTA, 1 mM DTT, 0.1 mg/ml BSA, 2% glycerol. Reactions were incubated at 37°C for 15 minutes to form open complexes, mixed with 1 mM of NCIN (NAD^+^, NADH, dpCoA, or FAD), 6 mCi [α-^32^P]-UTP (3000 Ci/mmol), 200 μM CTP, 200 μM UTP, and 200 μM GTP and incubated at 37°C for 10 minutes to allow for product formation. Reactions were stopped by addition of 1.5x stop solution (0.6 M Tris HCl pH 8.0, 18 mM EDTA, 0.1 mg/ml glycogen). Samples were extracted with acid phenol:chloroform and NCIN-capped products were recovered by ethanol precipitation.

NudC processing reactions were performed by adding 400 nM NudC to products resuspended in 10 μl of NudC reaction buffer (10 mM Tris HCl pH 8.0, 50 mM NaCl, 10 mM MgCl_2_, 1 mM DTT). Rai1 processing reactions were performed by adding 400 nM Rai1 to RNA products resuspended in 10 μl Rai1 reaction buffer [10 mM Tris-HCl (pH 7.5), 100 mM KOAc, 2 mM MgOAc, 1 mM MnCl_2_, 2 mM DTT]. Rpp processing reactions were performed by adding 10U Rpp to RNA products resuspended in 10 μl of Rpp reaction buffer [50 mM HEPES-KOH (pH 7.5), 100 mM NaCl, 1 mM EDTA, 0.1% β-mercaptoethanol, and 0.01% Triton X-100]. Reactions were incubated at 37°C for 30 minutes, stopped by addition 10 μl 2x RNA loading dye (95% deionized formamide, 18 mM EDTA, 0.25% SDS, xylene cyanol, bromophenol blue, amaranth), and separated by electrophoresis on 8% TBE-Urea polyacrylamide gels (UreaGel system, National Diagnostics). Radioactive products were detected by autoradiography using storage phosphor screens and a Typhoon 9400 variable mode imager (GE Life Science).

### CapZyme-Seq

#### Isolation of RNA products generated in vitro

A linear DNA fragment containing the *placCONS-N7* promoter library was used as a template for *in vitro* transcription assays. To generate this template, plasmid pMASTER-*lacCONS-N7* (Vvedenskaya et al., 2015) was diluted to ~10^9^ molecules/μl. 1μl of diluted DNA was amplified by emulsion PCR (ePCR) using a Micellula DNA Emulsion and Purification Kit in detergent-free Phusion HF reaction buffer containing 5 μg/ml BSA, 0.4 mM dNTPs, 0.5 μM Illumina RP1 primer, 0.5 μM Illumina RPI1 primer and 0.04 U/μl Phusion HF polymerase. ePCR reactions were performed with an initial denaturation step of 10 seconds at 95°C, amplification for 30 cycles (denaturation for 5 seconds at 95°C, annealing for 5 seconds at 60°C and extension for 15 seconds at 72°C), and a final extension for 5 minutes at 72°C. DNA was purified from emulsions according to the manufacturer’s recommendations, recovered by ethanol precipitation, and resuspended in nuclease-free water to a final concentration of ~1 μM.

*In vitro* transcription was performed by mixing 10 nM of template DNA with 50 nM RNAP holoenzyme in 50 mM Tris HCl (pH 8.0), 10 mM MgCl_2_, 0.01 mg/ml BSA, 100 mM KCl, 5% glycerol, 10 mM DTT, and 0.4U/μl RNase OUT. Reactions were incubated at 37°C for 10 minutes to form open complexes. A single round of transcription was initiated by addition of 100 μM ATP, 100 μM CTP, 100 μM UTP, 100 μM GTP, and 0.1 mg/ml heparin, or by addition of 100 μM ATP, 100 μM CTP, 100 μM UTP, 100 μM GTP, 2 mM NAD^+^, and 0.1 mg/ml heparin. Reactions were incubated at 37°C for 15 minutes and stopped by addition of 0.5M EDTA (pH 8) to a final concentration of 50 mM. Nucleic acids were recovered by ethanol precipitation and resuspended in 30 μl of nuclease-free water. Reactions performed in the absence or presence of NAD^+^ were performed in triplicate, and each replicate was analyzed by both CapZyme-Seq^NudC^ and CapZyme-Seq^Rai1^.

The resuspended nucleic acids were mixed with 30 μl of 2x RNA loading dye and separated by electrophoresis on 10% 7M urea slab gels (equilibrated and run in 1x TBE). The gel was stained with SYBR Gold nucleic acid gel stain, bands were visualized on a UV transilluminator, and RNA products ~100 nt in size were excised from the gel. The excised gel slice was crushed, 300 μl of 0.3 M NaCl in 1x TE buffer was added, and the mixture was incubated at 70°C for 10 minutes. Eluted RNAs were collected using a Spin-X column. After the first elution, the crushed gel fragments were collected and the elution procedure was repeated, nucleic acids were collected, pooled with the first elution, isolated by ethanol precipitation, and resuspended in 10 μl of RNase-free water.

#### Isolation of RNAs generated in vivo in E. coli

The pMASTER-*lac*CONS-N7 plasmid library is described in (Vvedenskaya et al., 2015) and contains the promoter cassette shown in Figure 1A fused to the tR2 terminator. RNA products generated from *lac*CONS promoter sequences that terminate at tR2 are ~100-nt in length. To analyze RNA products generated from the pMASTER-*lac*CONS-N7 library *in vivo*, three independent 50 ml cell cultures of DH10B-T1^R^ cells containing the pMASTER-*lac*CONS-N7 plasmid library [prepared as described in (Vvedenskaya et al., 2015)] were grown in LB media containing chloramphenicol (25 μg/ml) in a 250 ml DeLong flask shaken at 210 RPM at 37°C to mid-exponential phase (OD600 ~0.5). 2 ml aliquots of the cell suspensions were placed in 2 ml tubes and cells were collected by centrifugation (1 min, 21,000 x g at room temperature). Supernatants were removed and cell pellets were rapidly frozen on dry ice and stored at −80°C.

RNA was isolated from each cell pellet as in (Vvedenskaya et al., 2015). Cell pellets were resuspended in 600 μl of TRI Reagent solution. The suspensions were incubated at 70°C for 10 min and centrifuged (10 min, 21,000 x g) to remove insoluble material. The supernatant was transferred to a fresh tube, ethanol was added to a final concentration of 60.5%, and the mixture was applied to a Direct-zol spin column. DNase I treatment was performed on-column according to the manufacturer’s recommendations. RNA was eluted from the column using nuclease-free water that had been heated to 70°C (3 x 30 μl elutions; total volume of eluate = 90 μl). RNA was treated with 2U TURBO DNase at 37°C for 1 h to remove residual DNA. Samples were extracted with acid phenol:chloroform, RNA was recovered by ethanol precipitation and resuspended in RNase-free water. A MICROBExpress Kit was used to deplete rRNAs from 9 μg of the recovered RNA. The rRNA-depleted RNA was isolated by ethanol precipitation and resuspended in 30 μl of RNase-free water. pMASTER-lacCONS-N7 plasmid DNA was isolated from each of the three cultures using a Plasmid Mini-prep kit. Plasmid DNA isolated from these cultures was used as template in ePCR reactions to generate linear DNA products that were analyzed by high-throughput sequencing (DNA template libraries Vv828, Vv830, and Vv832, see below).

#### Enzymatic treatments of RNA products generated in vitro with Rpp, NudC, or Rai1

To convert 5′ triphosphate RNA to 5′ monophosphate RNA, products were mixed with 20U Rpp and 40U RNaseOUT in 1x Rpp reaction buffer in a 20 μl reaction volume.

To convert NAD^+^-capped RNA to 5′ monophosphate RNA, products were mixed with 1x NudC reaction buffer, 3.6 uM NudC, and 40U of RNaseOUT in a 20 μl reaction volume or with 1x Rai1 reaction buffer, 0.3 uM Rai1, and 40U RNaseOUT in a 20 μl reaction volume. In parallel, we added RNA products to each of the reaction buffers without addition of enzyme. Reactions were incubated at 37°C for 30 minutes, 20 μl of 2x RNA loading dye was added, and products were separated by electrophoresis on 10% 7M urea slab gels (equilibrated and run in 1x TBE). The gel was stained with SYBR Gold nucleic acid gel stain, bands were visualized on a UV transilluminator, and RNA products ~100 nt in size were excised from the gel. The excised gel slice was crushed, 300 μl of 0.3 M NaCl in 1x TE buffer was added, and the mixture was incubated at 70°C for 10 minutes. RNAs were collected using a Spin-X column. After the first elution, the crushed gel fragments were collected and the elution procedure was repeated, nucleic acids were collected, pooled with the first elution, isolated by ethanol precipitation, and resuspended in 10 μl of RNase-free water.

#### Enzymatic treatments of RNA products generated in vivo with Rpp or NudC

To convert 5′-triphosphate RNA to 5′-monophosphate RNA, 2 μg rRNA-depleted RNA were mixed with 20U Rpp and 40U RNaseOUT in 1x Rpp reaction buffer in a 30 μl reaction volume. In parallel, we added 2 μg rRNA-depleted RNA to 1x Rpp reaction buffer without addition of enzyme. Reactions were incubated at 37°C for 30 minutes. Samples were extracted with acid phenol:chloroform, RNA was recovered by ethanol precipitation, and resuspended in 10 μl RNase-free water.

Prior to treating total cellular RNA with NudC to convert NCIN-capped RNA to 5′-monophosphate RNA we first treated 2 μg of rRNA-depleted RNA with 2U CIP in the presence of 40U RNaseOUT in a 30 μl reaction volume at 37°C for 1h to remove 5′-terminal phosphates. RNA was extracted with acid phenol:chloroform, recovered by ethanol precipitation, and resuspended in 20 μl of RNase-free water. CIP-treated RNA was mixed with 3.75 μM NudC and 40U RNaseOUT in 1x NudC reaction buffer in a 30 μl reaction volume. In parallel, CIP-treated RNA was incubated in 1x NudC reaction buffer without NudC. Reactions were incubated at 37°C for 30 minutes, 30 μl of 2x RNA loading dye was added, and products were separated by electrophoresis on 10% 7M urea slab gels (equilibrated and run in 1x TBE). The gel was stained with SYBR Gold nucleic acid gel stain, RNA was visualized on a UV transilluminator, and excised from the gel. The excised gel slice was crushed, 400 μl of 0.3 M NaCl in 1x TE buffer was added, and the mixture was incubated at 70°C for 10 minutes. Eluted RNAs were separated from crushed gel fragments using a Spin-X column. After the first elution, the crushed gel fragments were collected and the elution procedure was repeated, nucleic acids were collected, pooled with the first elution, isolated by ethanol precipitation, and resuspended in 10 μl of RNase-free water.

#### 5′ adaptor ligation

RNA products (in 10 μl of nuclease-free water) were combined with PEG 8000 (10% final concentration), 1 mM ATP, 40U RNaseOUT, 1x T4 RNA ligase buffer, and 10 U T4 RNA ligase 1, in 30 μl reaction volume. 0.3 μM barcoded 5′ adaptor oligo was added to *in vitro* generated RNAs, and 1 μM barcoded 5′ adaptor oligo was added to *in vivo* generated RNAs, respectively. Reactions were incubated at 16°C for 16 h.

To enable quantitative comparisons between samples treated with Rpp, samples treated with NCIN-processing enzymes, samples incubated in Rpp reaction buffer (“mock Rpp treatment”), and samples incubated in NCIN-processing enzyme reaction buffer (“mock NudC treatment” or “mock Rai1 treatment”), we performed the 5′ adaptor ligation step using barcoded 5′-adaptor oligonucleotides. For libraries prepared from RNA generated *in vitro* or *in vivo*, oligo i105 was used in ligation reactions performed with products isolated after Rpp treatment, oligo i106 was used in ligation reactions performed with products isolated after mock Rpp treatment, oligo i107 was used in ligation reactions performed with products isolated after NudC treatment (for CapZyme-Seq^NudC^) or Rai1 treatment (for CapZyme-Seq^Rai1^), and oligo i108 was used in ligation reactions performed with products isolated after mock NudC treatment or mock Rai1 treatment.

The ligation reactions were stopped by adding 30 μl of 2x RNA loading dye and heated at 95°C for 5 minutes. Each set of four adaptor ligation reactions were combined, mixed with an equal volume of 2x RNA loading dye, and separated by electrophoresis on 10% 7M urea slab gels (equilibrated and run in 1x TBE). Gels were incubated with SYBR Gold nucleic acid gel stain, and bands were visualized with UV transillumination. For *in vitro* generated RNAs, species ~150 nt in length were recovered from the gel (procedure as above) and resuspended in 10 μl of nuclease-free water. For *in vivo* generated RNAs, species migrating above the 5′-adapter oligo were recovered from the gel (procedure as above) and resuspended in 50 μl of nuclease-free water.

#### First strand cDNA synthesis: analysis of RNAs generated from the lacCONS-N7 promoter library in vitro

5′-adaptor-ligated products (in 10 μl of nuclease-free water) were mixed with 1.5 μM s128A oligonucleotide, incubated at 65°C for 5 minutes, then cooled to 4°C. To this mixture was added 9.7 μl of a solution containing 4 μl of 5x First-Strand buffer, 1 μl of 10 mM dNTP mix, 1 μl of 100 mM DTT, 1 μl (40U) RNaseOUT, 1 μl (200U) of SuperScript III Reverse Transcriptase and 1.7 μl of nuclease-free water. Reactions were incubated at 25°C for 5 minutes, 55°C for 60 minutes, 70°C for 15 minutes, then cooled to 25°C. 10U of RNase H (Life Technologies) was added, the reactions were incubated at 37°C for 20 minutes and 20 μl of 2x RNA loading dye was added. Nucleic acids were separated by electrophoresis on 10% 7M urea slab gels (equilibrated and run in 1x TBE). Gels were incubated with SYBR Gold nucleic acid gel stain, bands were visualized with UV transillumination, and species ~80 to ~150 nt in length were recovered from the gel (procedure as above) and resuspended in 20 μl of nuclease-free water.

#### First strand cDNA synthesis: analysis of RNAs generated in vivo

5′-adaptor-ligated products were divided into two equal portions (each in 25 μl of nuclease-free water). One portion was mixed with 0.5 μl of 100 μM s128A oligonucleotide to enable analysis of RNA products generated from the lacCONS-N7 promoter library, while the other portion was mixed with 0.5 μl of 100 μM sRNA oligo pool (a mixture of 77 oligonucleotides each having a 3′-end sequence complementary to positions +50 to +30 of one of 77 annotated sRNAs in *E. coli*; see Key Resources Table). The mixtures were incubated at 65°C for 5 min, then cooled to 4°C. To these mixtures was added 24.5 μl of a solution containing 10 μl 5x First-Strand buffer 2.5 μl 10 mM dNTP mix, 2.5 μl 100 mM DTT, 2.5 μl (100U) RNaseOUT, 2.5 μl (500U) SuperScript III Reverse Transcriptase and 4.5 μl of nuclease-free water. Reactions were incubated at 25°C for 5 min, 55°C for 60 min, 70°C for 15 min, then cooled to 25°C. 20U RNase H was added, the reactions were incubated at 37°C for 20 min and 50 μl of 2x RNA loading dye was added. Nucleic acids were then separated by electrophoresis on 10% 7M urea slab gels. To isolate cDNAs derived from the *lac*CONS-N7 promoter library, species ~80 to ~150 nt in length were recovered from the gel (procedure as above) and resuspended in 20 μl of nuclease-free water. To isolate cDNAs derived from sRNAs, species ~80 to ~225 nt in length were recovered from the gel and resuspended 20 μl of nuclease-free water.

#### cDNA amplification

cDNA derived from RNA products generated *in vitro* were diluted 10-fold with nuclease-free water. cDNAs derived from RNA products generated *in vivo* were diluted 20-to 30-fold with nuclease-free water. 2μl of the diluted cDNA solution was used as a template for ePCR reactions containing Illumina index primers using a Micellula DNA Emulsion and Purification Kit (20 cycles; conditions as above). The emulsion was broken and DNA was purified according to the manufacturer’s recommendations. DNA was recovered by ethanol precipitation and resuspended in 20 μl of nuclease-free water.

#### High-throughput sequencing

Barcoded libraries were pooled and sequenced on an Illumina NextSeq platform in high-output mode using custom primer s1115.

#### Sample serial numbers for CapZyme-Seq^Rail^ in vitro

Samples Vv1225, Vv1229, Vv1230, and Vv1231 are cDNA derived from RNA products generated *in vitro* in the absence of NAD^+^. Vv1225 and Vv1229 are cDNA generated from the same RNA products, and thus were considered as a single replicate. Samples Vv1227, Vv1232, Vv1233, and Vv1234 are cDNA derived from RNA products generated *in vitro* in the presence of NAD^+^. Samples Vv1227 and Vv1232 are cDNA generated from the same RNA products, and thus were considered as a single replicate.

#### Sample serial numbers for CapZyme-Seq^NudC^ in vitro

Samples Vv1226, Vv1235, Vv1236, and Vv1237 are cDNA derived from RNA products generated *in vitro* in the absence of NAD^+^. Samples Vv1226 and Vv1235 are cDNA generated from the same RNA products and thus were considered as a single replicate. Samples Vv1228, Vv1238, Vv1239, and Vv1240 are cDNA derived from RNA products generated *in vitro* in the presence of NAD^+^. Samples Vv1228 and Vv1238 are cDNA libraries generated from the same RNA products, and thus were considered as a single replicate Sample Vv1168 is the DNA template used in the MASTER-*lac*CONS-N7 *in vitro* reactions.

#### Sample serial numbers for CapZyme-Seq^NudC^ in vivo

Samples Vv1273, Vv1275, and Vv1277 are cDNA derived from RNA products generated *in vivo* from the *lac*CONS-N7 promoter library. Samples Vv1274, Vv1276, and Vv1278 are cDNA derived from sRNAs products present in total cellular RNA isolated from *E. coli*. Samples Vv828, Vv830, and Vv832 are DNA products generated in ePCR reactions using pMASTER-lacCONS-N7 plasmid DNA isolated from cells.

### CapZyme-Seq data analysis

Source code and documentation for CapZyme-Seq analysis are provided at: https://github.com/NickelsLabRutgers

#### Transcribed-region barcode identification for MASTER-lacCONS-N7 experiments

The DNA template used for *in vitro* transcription reactions was analyzed by high-throughput sequencing (sample serial number Vv1168) to identify transcribed-region barcodes, and to assign these barcodes to individual placCONS template sequences as described previously (Vvedenskaya et al., 2015).

For *in vivo* MASTER-lacCONS-N7 experiments, DNA products generated in ePCR reactions using pMASTER-*lac*CONS-N7 plasmid DNA isolated from cells were analyzed by high-throughput sequencing (sample serial numbers Vv828, Vv830, and Vv832) to identify transcribed-region barcodes, and to assign these barcodes to individual p*lac*CONS template sequences as described previously (Vvedenskaya et al., 2015). In particular, analysis of sample Vv828 was used to assign transcribed-region barcodes for analysis of sample Vv1273, analysis of sample Vv830 was used to assign transcribed-region barcodes for analysis of sample Vv1275, and analysis of sample Vv832 was used to assign transcribed-region barcodes for analysis of sample Vv1277.

#### Analysis of sequencing reads derived from cDNA

All cDNA libraries were generated from the same input RNA that had been split into four portions and subjected to two distinct 5′ processing reactions (Rpp and NudC or Rai1) and two distinct control reactions (mock Rpp treatment and mock NudC or Rai1 treatment). To distinguish cDNA derived from each of the four reactions, RNA from each reaction was ligated to a 5′-adaptor oligonucleotide containing a unique 4-nt barcode sequence (i105, i106, i107, or i108; see above). Each 5′-adaptor oligonucleotide also contains 11 nt of random sequence at the 3′ end that improves ligation efficiency (by reducing sequence-dependent effects), and that marks individual RNA products with an 11-nt sequence tag that is used to reduce effects of PCR amplification bias. Thus, sequencing reads having identical 11-nt sequence tags and identical cDNA insert sequences are counted as a single read count during the data analysis.

Due to the presence of the 4-nt barcode sequence and 11-nt sequence tag at the 3′ end of the 5′-adaptor oligonucleotide, the first four bases of each read provide the sequence of the 4-nt barcode, the next 11 bases provide the sequence of the 11-nt sequence tag, and the 16^th^ base provides the sequence of the RNA 5′ end from which the cDNA was generated.

#### Analysis of sequencing reads derived from cDNA: MASTER-lacCONS-N7 analysis in vitro

For analysis of cDNA libraries generated from RNA products produced from the MASTER-*lac*CONS-N7 library *in vitro* the 15-base transcribed-region barcode was identified as described in (Vvedenskaya et al., 2015) and used to associate reads derived from RNA transcripts with their template of origin. We considered only RNA 5′-end-sequences that could be aligned to the sequence of their template of origin with no mismatches (Vvedenskaya et al., 2015). These reads were associated with one of the four reaction conditions based on the identity of the 4-nt barcode sequence.

For each of the ~16,000 sequence variants, we determined the number of reads emanating from each position of the N7 region for samples treated with Rpp (#Rpp), samples treated with NCIN-processing enzymes (#NudC or #Rai1), samples subjected to mock Rpp treatment (#Rpp^mock^), and samples subjected to mock NCIN-processing enzyme treatment (NudC^mock^ or #Rai1^mock^). From these values, we calculated #ppp and #NCIN, where #ppp = (#Rpp - #Rpp^mock^), and #NCIN = (#NudC - #NudC^mock^) for CapZyme-Seq^NudC^ and #NCIN = (#Rai1 - #Rai1^mock^) for CapZyme-Seq^Rai1^. Negative values for #ppp or #NCIN were replaced with a value of “0”.

Analysis of reactions performed in the absence of NAD^+^ revealed activity of NudC and Rai1 on 5′-triphosphate RNA. By comparison with analysis of reactions performed in the absence of NAD^+^ with Rpp we estimate, on average, NudC converted ~19% of 5′-triphosphate RNA to 5′-monophosphate RNA, and Rai1 converted ~3.5% of 5′-triphosphate RNA to 5′-monophosphate RNA. Therefore, to account for the conversion of 5′-triphosphate RNA to 5′-monophosphate RNA by NudC or Rai1 we used values of #ppp and #NCIN obtained in reactions performed in the absence of NAD^+^ to calculate a correction factor (*cf*), where *cf* = (#NCIN / #ppp), to apply to the analysis of reactions performed in the presence of NAD^+^.

To analyze results of reactions performed in the presence of NAD^+^ we used the value of *cf*, the values for #ppp, and the value for #NCIN to calculate a “background corrected” value of #NCIN (#NCIN^Bkd_Cor^), where #NCIN^Bkd_Cor^ = [#NCIN - (*cf* x #ppp)]. Next, using the value of #NCIN^Bkd_^ and #ppp we calculated a value of “percent capping,” where percent capping = 100% x [#NCIN^Bkd_Cor^ / (#NCIN^Bkd_Cor^ + #ppp)]. (Note that values of (#NCIN^Bkd_Cor^ + #ppp) of 0 were replaced by “1” prior to calculating percent capping.)

For results of experiments performed *in vitro* we calculated a value for “capping efficiency,” where capping efficiency = [percent capping / (100 - percent capping)] / 20. (Note that the value of 20 corresponds to the [NAD^+^]/[ATP] for the *in vitro* transcription experiments performed in this work.

Results of Figure 2C represent a plot of the mean percent capping value (n=3; replicates of CapZyme^NudC^ or CapZyme^Rai1^) for template sequences with (#NCIN^Bkd_Cor^ + #ppp) ≥ 50.

Results of Figure 3B represent the mean TSS value (n=3; replicates of CapZyme^NudC^ or CapZyme^Rai1^) and mean %TSS values (n=6; replicates of CapZyme^NudC^ and CapZyme^Rai1^) for template sequences with #NCIN^Bkd^−^Cor^ ≥ 50. Results of Figure 3C and Figure S1B represent the mean TSS value (n=3; replicates of CapZyme^NudC^ or CapZyme^Rai1^) and mean %TSS values (n=6; replicates of CapZyme^NudC^ and CapZyme^Rai1^) for template sequences with #ppp ≥ 50. Mean TSS and %TSS were calculated as described (Vvedenskaya et al., 2015) with formulas given below.

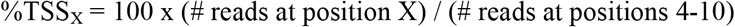

mean TSS = [(4 x %TSS at position 4) + (5 x %TSS at position 5) + (6 x %TSS at position 6) + (7 x %TSS at position 7) + (8 x %TSS at position 8) + (9 x %TSS at position 9) + (10 x %TSS at position 10)] / 100

Results of Figures 4 and 5B represent relative capping efficiency values calculated as described in Figure S2 for the position 7 bp downstream of the promoter -10 element for template sequences with (#NCIN^Bkd_Cor^ at position 7 + #ppp at position 7) ≥ 25 in each of the three CapZyme-Seq^NudC^ replicates and in each of the three CapZyme-Seq^Rai1^ replicates. Relative capping efficiency at each position was determined for groups of four promoter sequences, “quartets,” having A, G, C, or T, along with sequences identical at each other position. The relative capping efficiency of each A_+1_ promoter sequence, Y, at each position, X, is calculated by dividing the capping efficiency of Y by the mean capping efficiency of Y’s quartet at position X. For each promoter sequence, the mean capping efficiency (n=6; replicates of CapZyme^NudC^ and CapZyme^Rai1^) was used to calculate relative capping efficiency at each position. For results of Figure 5C, the relative capping efficiency of the consensus and anti-consensus promoter sequences were calculated by dividing the capping efficiency of each consensus or anti-consensus promoter sequence by the mean capping efficiency of all A_+1_ promoter sequences.

#### Analysis of sequencing reads derived from cDNA: MASTER-lacCONS-N7 analysis in vivo

For analysis of cDNA libraries generated from RNA products produced from the MASTER-*lac*CONS-N7 library *in vivo* the 15-base transcribed-region barcode was identified as described in (Vvedenskaya et al., 2015) and used to associate reads derived from RNA transcripts with their template of origin. As with RNAs generated *in vitro*, we also considered only RNA 5′-end-sequences that could be aligned to the sequence of their template of origin with no mismatches (Vvedenskaya et al., 2015). These reads were associated with one of the four reaction conditions based on the identity of the 4-nt barcode sequence.

For each of the ~16,000 sequence variants, we determined the number of reads emanating from each position of the N7 region for samples treated with Rpp (#Rpp), samples subjected to mock Rpp treatment (#Rpp^mock^), samples subjected to mock NudC treatment (#NudC), and samples subjected to mock NudC treatment (NudC^mock^). Using the values for #Rpp, #Rpp^mock^, #NudC and #NudC^mock^ we calculated values of #ppp and #NCIN as described above. To avoid complications due to the background activity of NudC on 5′-triphosphate RNA we treated RNA generated *in vivo* with CIP to remove 5′-terminal phosphates prior to treatment with NudC. In contrast to the analysis of RNA produced *in vitro*, removal of 5′-terminal phosphates by CIP prior to NudC treatment enabled us to directly use the value of #NCIN to calculate percent capping for RNA produced *in vivo*.

To calculate percent capping *in vivo* we used the value for #Rpp instead of that for #ppp in order to include 5′-monophosphate RNA in our analysis. Thus, we calculated a value of percent capping *in vivo* using the formula: percent capping = 100% x [#NCIN / (#NCIN + #Rpp)].

For results shown in Figures 7A-D, analysis was done using read count sums of the three independent CapZyme-Seq^NudC^ replicates. Results of Figure 7A represent a plot of the mean percent capping values for template sequences with (#NCIN + #Rpp) ≥ 50. Results of Figure 7B (left) and Figure S4A (left) represent the mean TSS value and mean %TSS values calculated using the sum of #NCIN of all template sequences. Results of Figure 7B (right) and Figure S4A (right) represent the mean TSS value and mean %TSS values calculated using the sum of #ppp of all template sequences. %TSS and the mean TSS were calculated as described above.

Results of Figures 7C and S4C,D represent relative difference in percent capping values calculated using the formula in Figure S4B for the position 7 bp downstream of the promoter −10 element for template sequences with (#NCIN at position 7 + #ppp at position 7) ≥ 25. Relative difference in percent capping at each position was determined for groups of four promoter sequences, “quartets,” having A, G, C, or T, along with sequences identical at each other position. The relative difference in percent capping of each A+1 promoter sequence, Y, at each position, X, is calculated by subtracting the percent capping of Y by the mean percent capping of Y’s quartet at position X. For results of Figure 7D, percent capping values of the consensus and anti-consensus promoter sequences are shown.

#### Analysis of sequencing reads derived from cDNA: sRNA analysis

Reads were associated with one of the four reaction conditions based on the identity of the 4-nt barcode sequence, the next 11 bases provide the sequence of the 11-nt sequence tag, and the 16^th^ base provides the sequence of the RNA 5′ end from which the cDNA was generated. The first 20 bases of the RNA 5′-end-sequence of each sequencing read was mapped exactly to the sequences from 50 bp upstream to 50 bp downstream of the annotated 5′-end position of each sRNA. Read counts derived from samples treated with Rpp (#Rpp), samples subjected to mock Rpp treatment (#Rpp^mock^), samples subjected to NudC treatment (#NudC), and the number of reads for samples subjected to mock NudC treatment (NudC^mock^). Using the values for #Rpp, #Rpp^mock^, #NudC and #NudC^mock^ we calculated values of #ppp and #NCIN as described above. As mentioned above, removal of 5′-terminal phosphates by CIP prior to NudC treatment enabled us to directly use the value of #NCIN to calculate percent capping for sRNA produced *in vivo*.

We first used values for #Rpp and #Rpp^mock^ to identify positions with the highest value of #Rpp for each sRNA. Next, for these positions we identified those representing primary TSS where the value of #Rpp was at least two times greater than #Rpp^mock^. In this manner, we identified 16 primary TSS positions where the base pair is A:T and the sum of #Rpp and #NCIN is greater than 100 in each of the three replicates (Figure 7E). For these primary TSS, we calculated a value of percent capping using the formula: percent capping = 100% x [#NCIN / (#NCIN + #Rpp)]. Values reported in Figure 7E are the mean of three replicates.

#### Data deposition

Raw reads have been deposited in the NIH/NCBI Sequence Read Archive under the study accession number PRJNA411835.

### Single-template gel analysis of NCIN capping with NAD^+^ (Bird et al., 2017)

10 nM of linear template was mixed with 50 nM RNAP holoenzyme in transcription buffer and incubated at 37°C for 15 minutes to form open complexes. 1 mM NAD^+^ (Roche), and increasing concentrations of ATP (GE Life Science) (0, 10, 20, 50, 100, 200, 500 μM) were added along with 200 nM of non-radiolabeled extending nucleotide (CTP, UTP or GTP; GE Life Science) plus 6 mCi of radiolabeled extending nucleotide ([α^32^P]-CTP, [α^32^P]-UTP, or [α^32^P]-GTP; Perkin Elmer; 3000 Ci/mmol). Upon addition of nucleotides, reactions were incubated at 37°C for 10 minutes to allow for product formation. Reactions were stopped by addition of an equal volume of gel loading buffer [90% formamide, 100 mM Tris-HCl pH 8.0, 18 mM EDTA, 0.025% xylene cyanol, 0.025% bromophenol blue].

Samples were run on 20% TBE-Urea polyacrylamide gels. Bands were quantified using ImageQuant software. Observed values of NAD^+^pC / (pppApC + NAD^+^pC) were plotted vs. [NAD^+^]/[ATP] on semi-log plot (Sigmaplot). Non-linear regression was used to fit the data to:

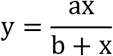

where y is NAD^+^pC / (pppApC + NAD^+^pC), x is [NAD^+^]/[ATP], and a and b are regression parameters. The resulting fit yields the value of x for which y = 0.5. The relative efficiency (k_cat_/K_M_, NAD^+^) / (k_cat_/K_M_, ATP) is equal to 1/x.

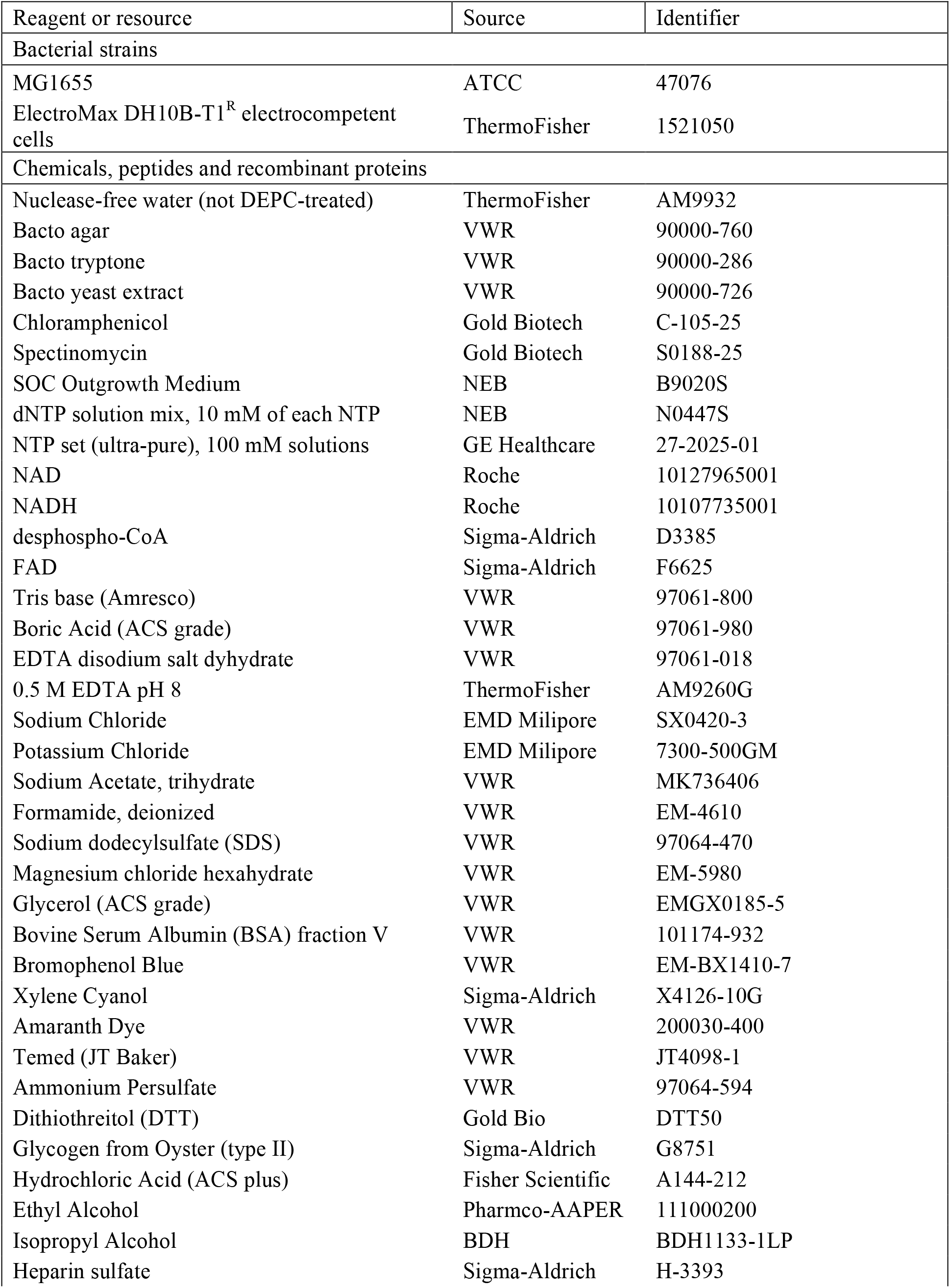
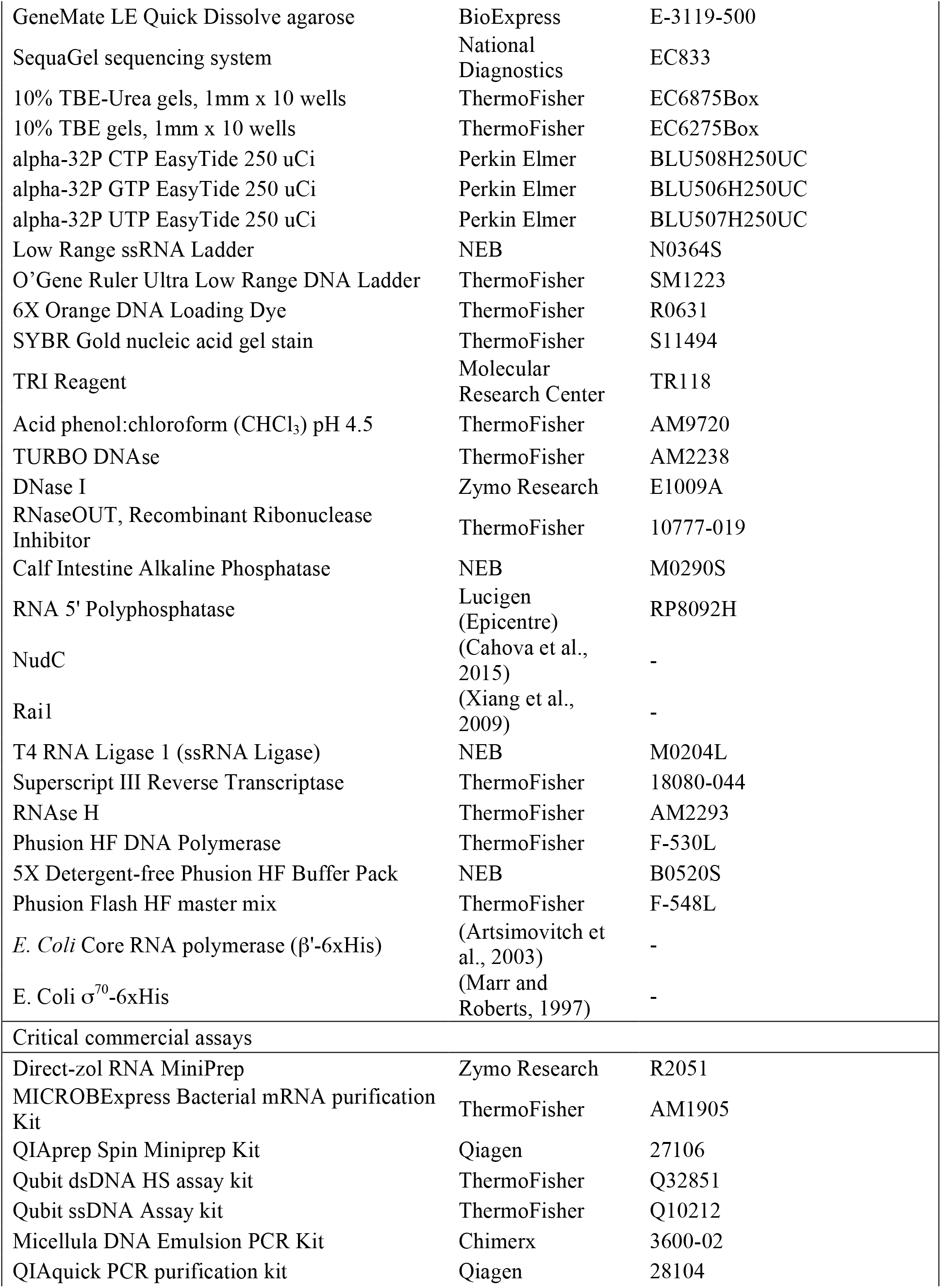
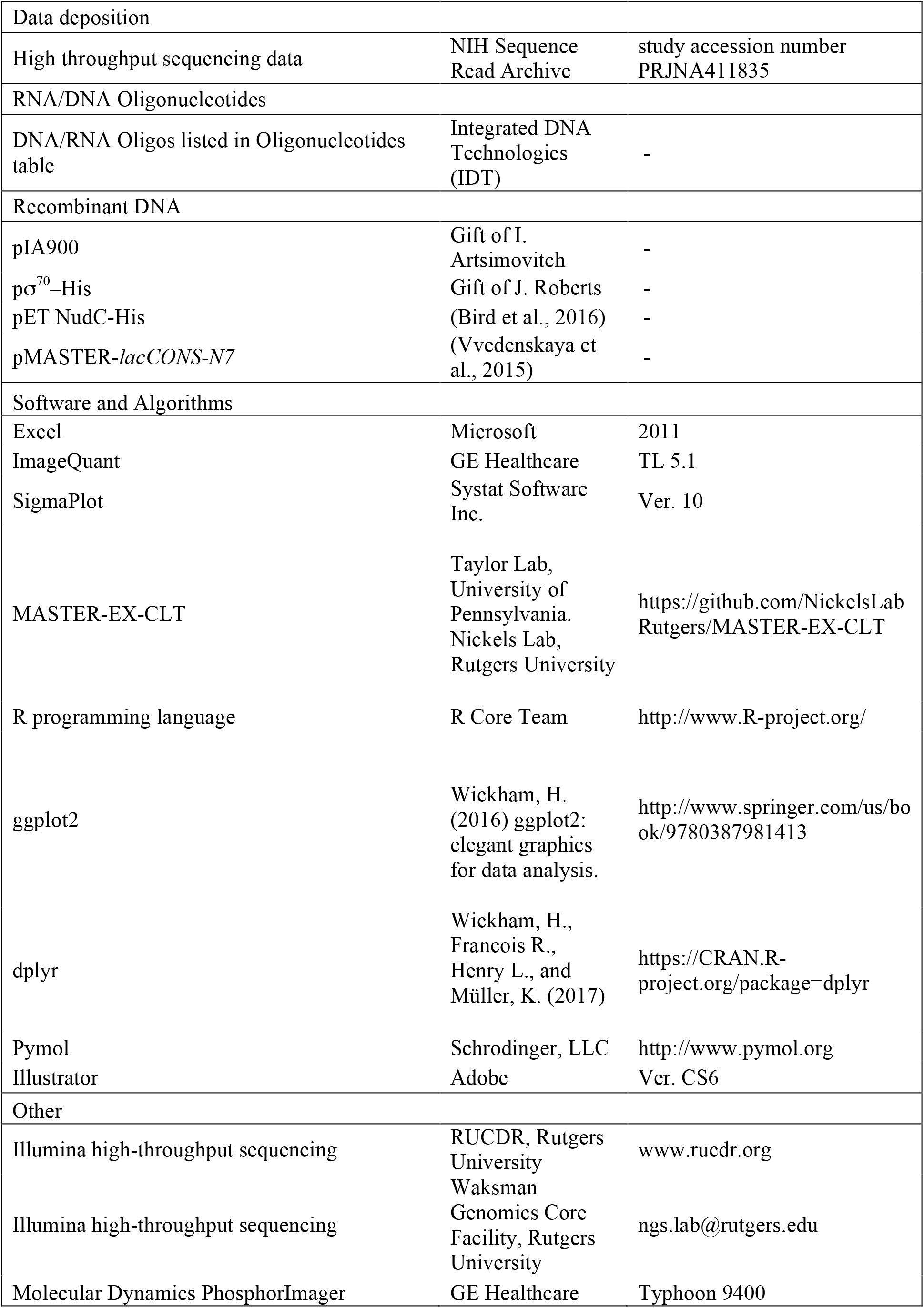
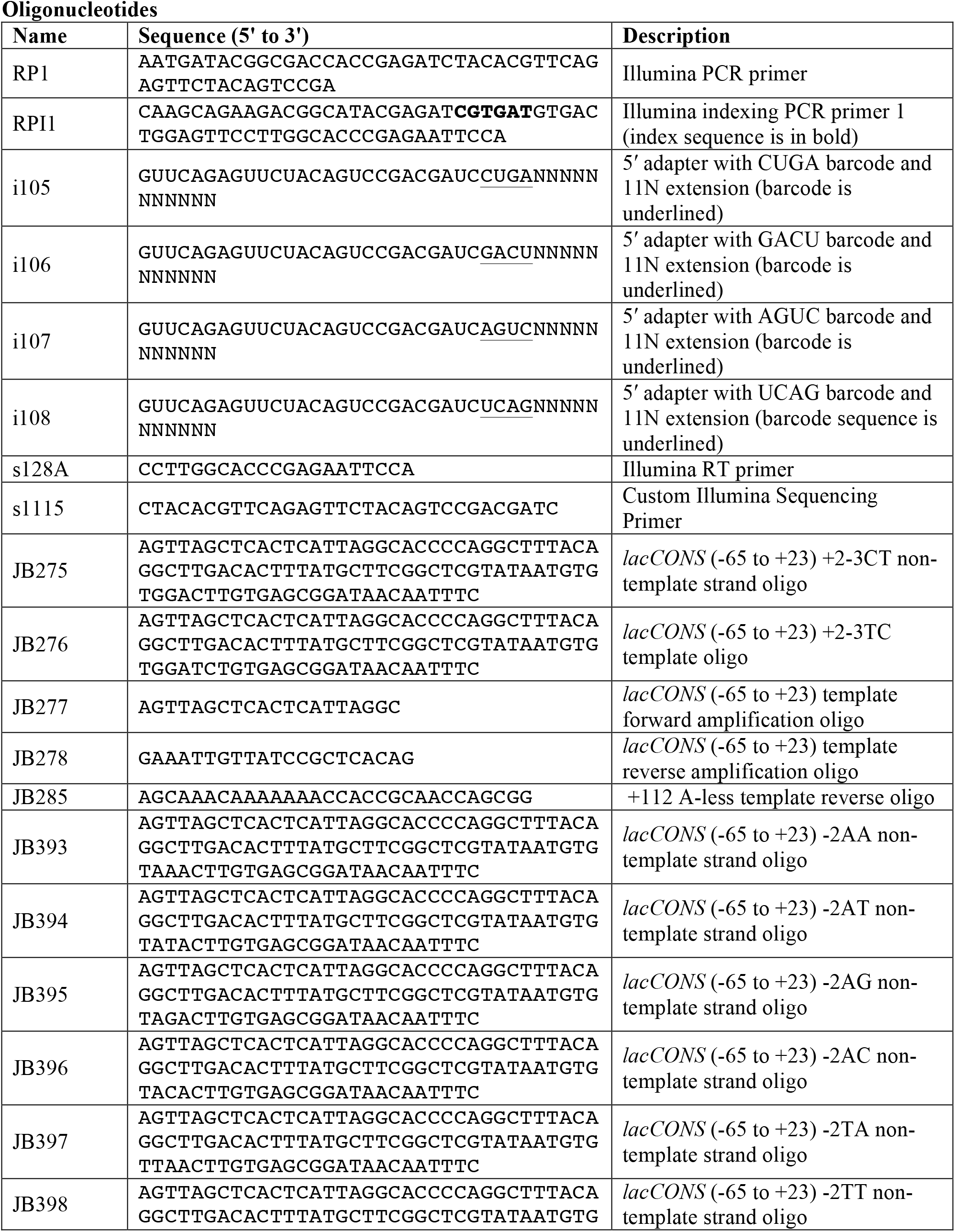
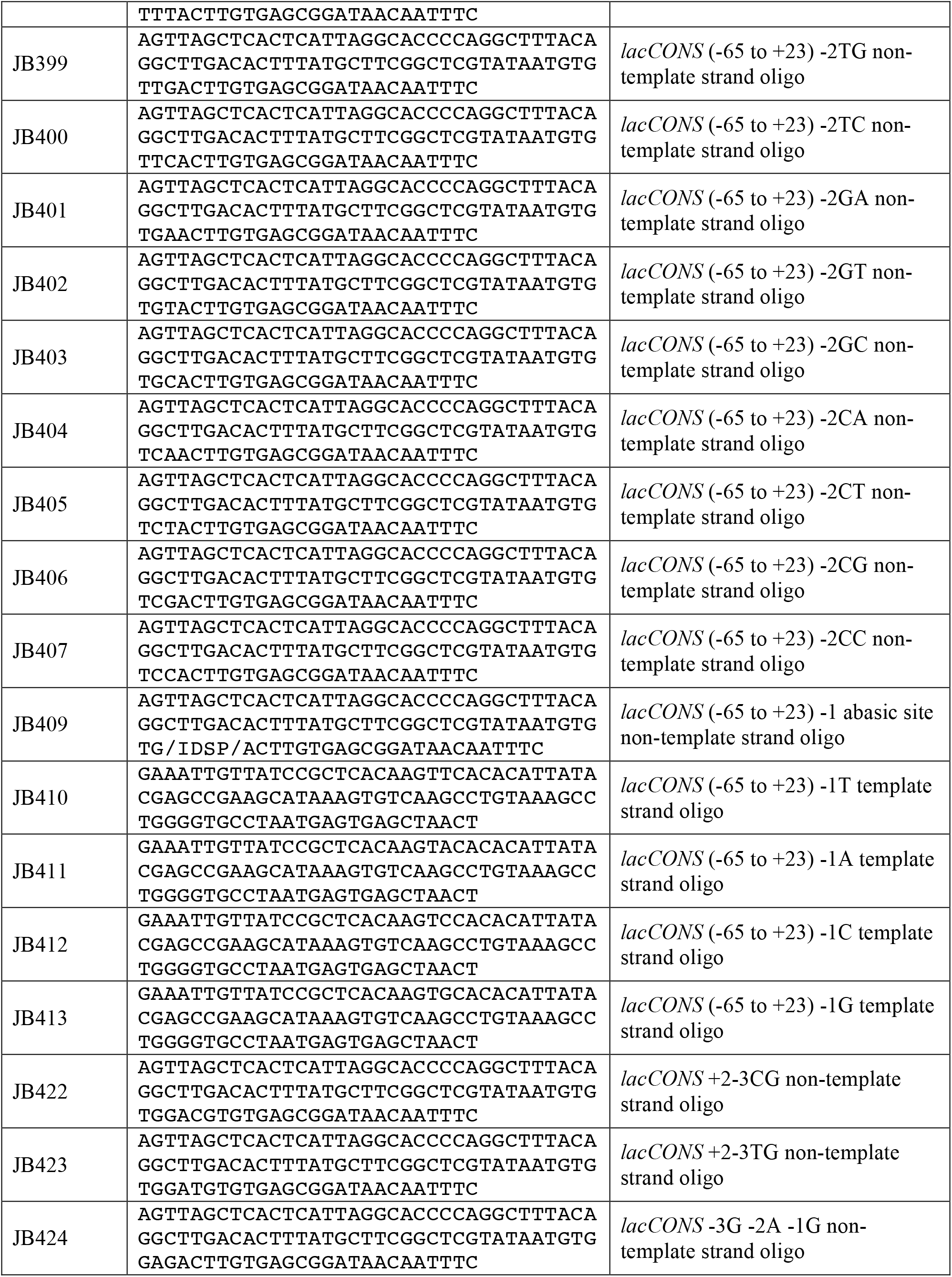
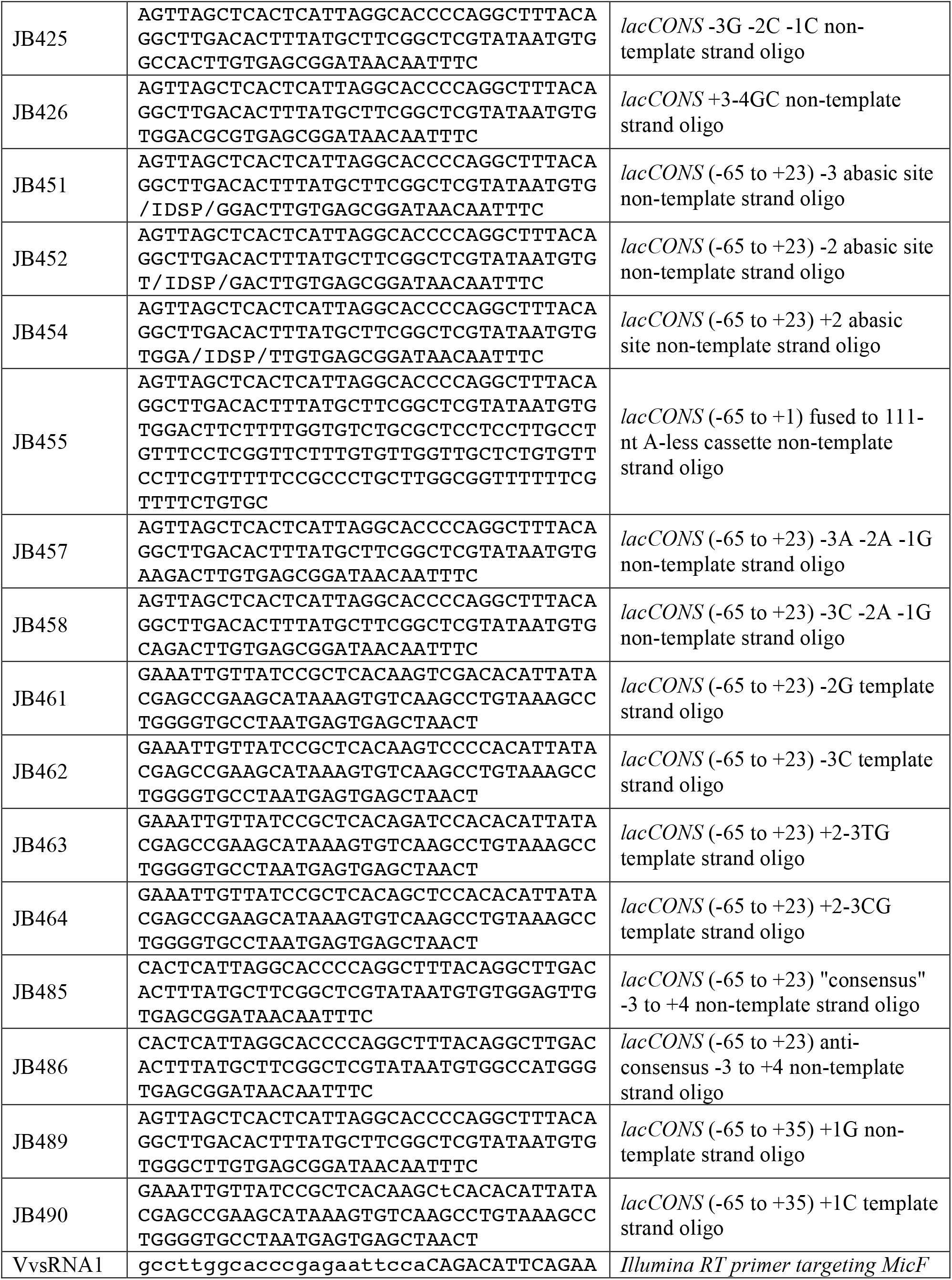
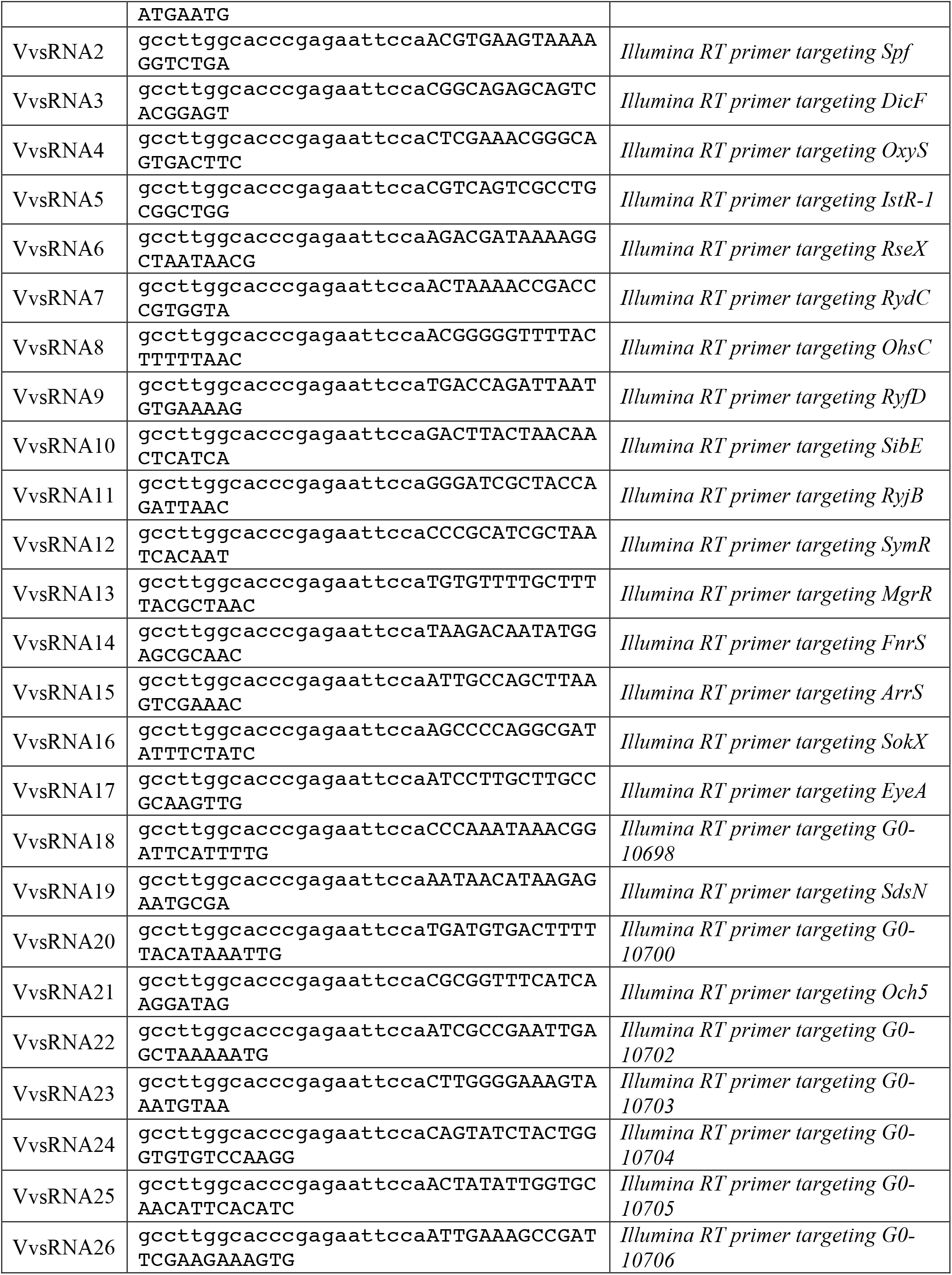
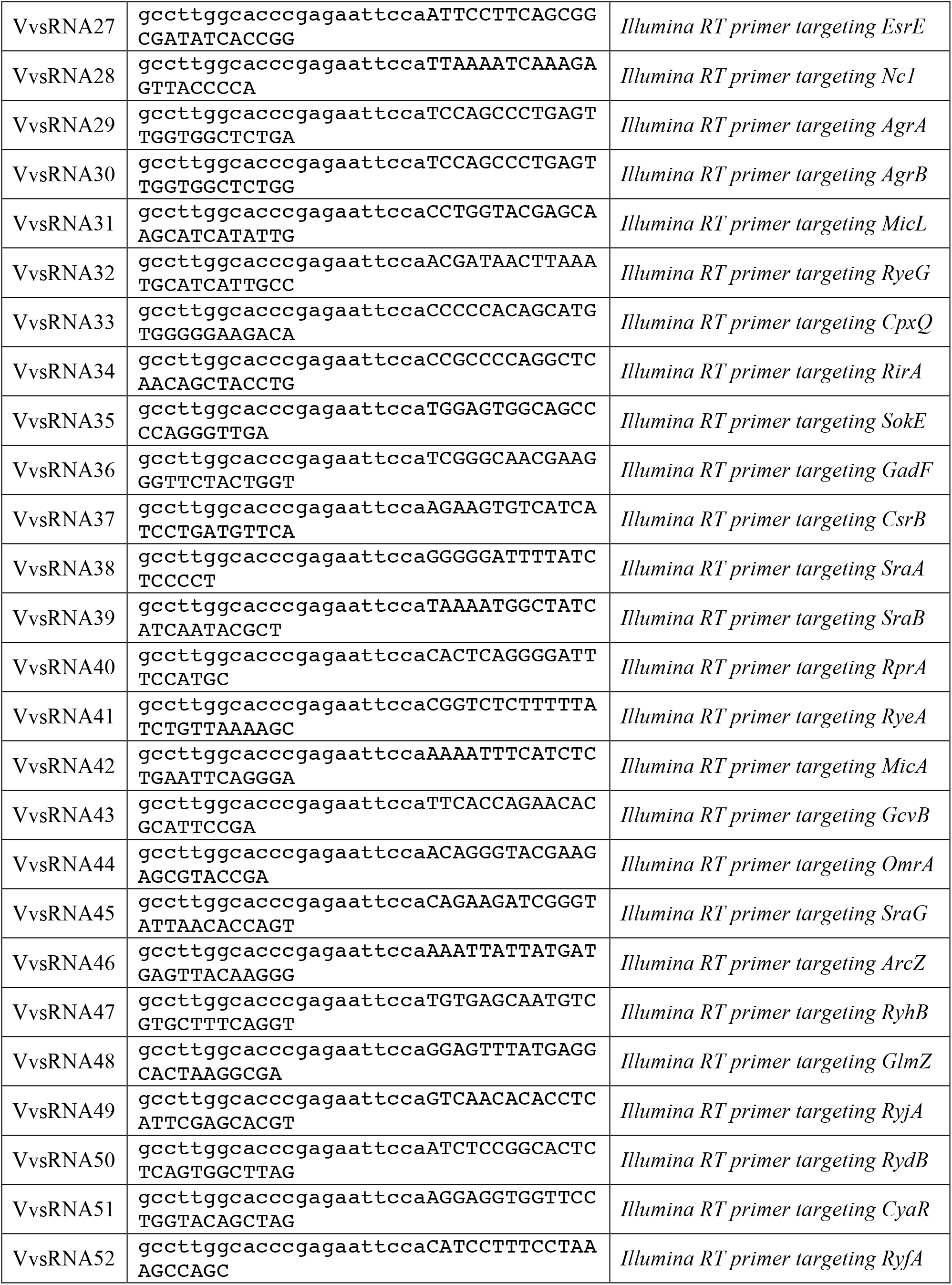
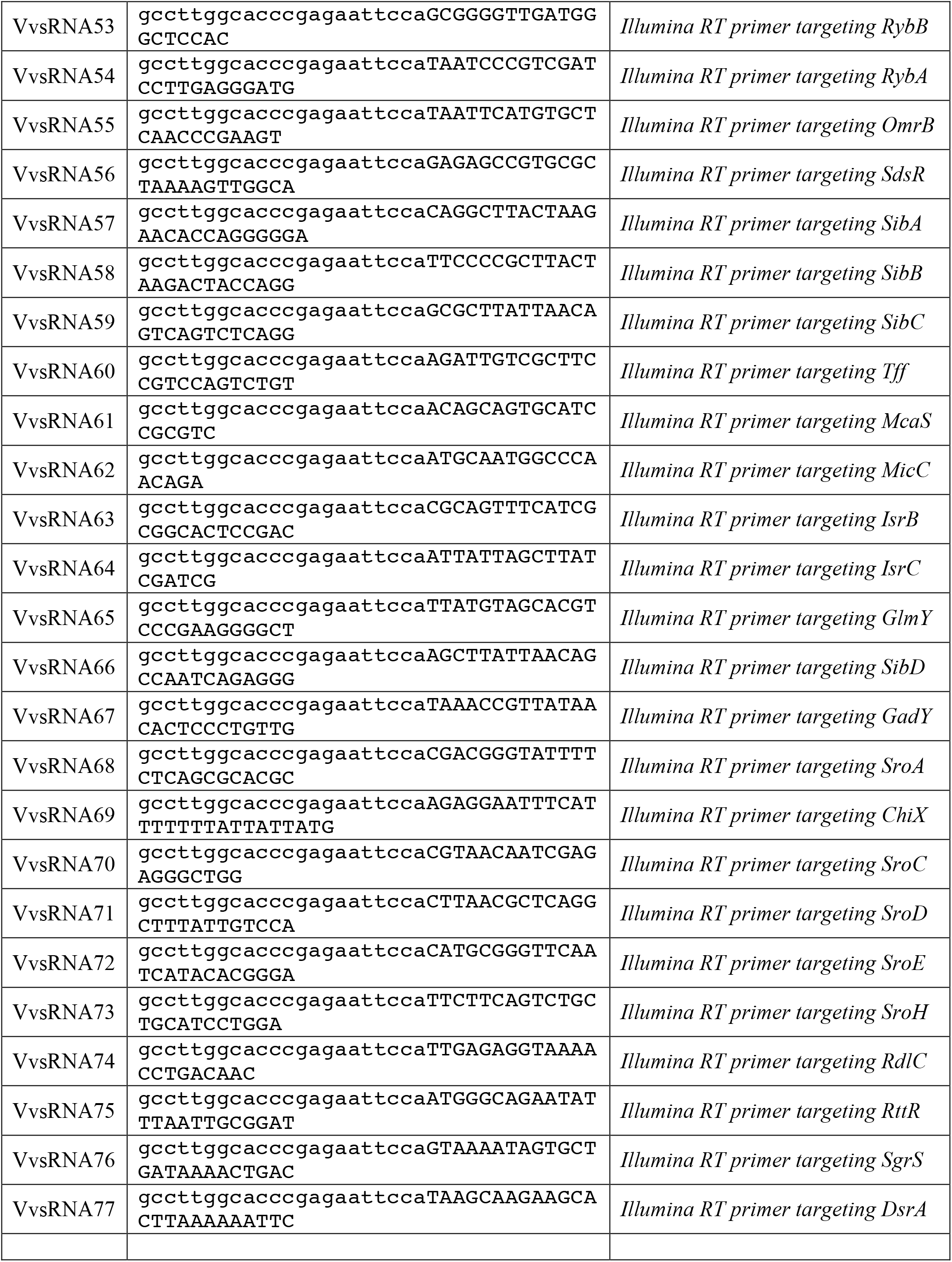
Key Resources Table

